# Modulation of the RNAse P/MRP complex and mitochondrial ribosome enhances cytosolic ribosome coordination and sustains longevity

**DOI:** 10.64898/2026.02.23.707525

**Authors:** Victoria E Martinez-Miguel, Till Popkes-Van Oepen, Tabrez Syed Shamsh, Francisco Lopes, Youngjun Park, Luca Sophie Jeromin, Edward Anderton, Wenming Huang, Gordon J Lithgow, Andreas Beyer, Adam Antebi

## Abstract

Aging is accompanied by a progressive decline in protein synthesis and ribosome abundance, yet paradoxically, genetic or pharmacological attenuation of translation extends lifespan across species. Whether the age-associated decline in translation is adaptive or reflects a pathological failure of ribosome homeostasis remains unclear. Here, we show that shortened lifespan is driven by dysregulated ribosome biogenesis (RiBi) and impaired ribosome assembly. Using *Caenorhabditis elegans ncl-1* loss-of-function mutants as a model of accelerated aging, we find that nucleolar enlargement coincides with decoupling of precursor and mature rRNA, ribosomal protein (RP) transcripts, and protein abundance, loss of RP stoichiometry, defective ribosomal subunit joining, and compromised proteostasis. Strikingly, lifespan can be restored downstream of the nucleolus by targeting either the RNAse P/MRP complex or the mitochondrial ribosome. These interventions rebalance mature rRNA and RP abundance, improve ribosomal assembly, and reduce protein aggregation despite persistent nucleolar enlargement and elevated pre-rRNA levels. Our findings identify accelerated age-associated ribosome dysfunction as a qualitative failure of ribosomal biogenesis, and demonstrate that restoring ribosomal homeostasis is sufficient to improve proteostasis and extend lifespan.

## Introduction

Protein synthesis is a highly regulated and energetically demanding process essential to healthy cellular function, and its dysregulation is a conserved feature of aging. Across taxa, aging is associated with a progressive decline in both global protein synthesis (Blazejowski & Webster, 1983; Dwyer et al., 1980; Fando et al., 1980; Layman et al., 1976; Motizuki & Tsurugi, 1992; Stein et al., 2022; Ward & Richardson, 1991; Webster & Webster, 1979) and ribosomal protein (RP) abundance (Walther et al., 2015; Ubaida-Mohien et al., 2019; Llewellyn et al., 2023; Lu et al., 2023; H. Zhu et al., 2023; Anisimova et al., 2020; Di Fraia et al., 2025; Brown et al., 2018; Dhondt et al., 2017).

Paradoxically, many lifespan-extending interventions downregulate global protein synthesis, and genetic or pharmacological attenuation of translation is sufficient to extend lifespan in multiple organisms (Bitto et al., 2016; Chen et al., 2007; Chiocchetti et al., 2007; Curran & Ruvkun, 2007; Demontis & Perrimon, 2010; Hansen et al., 2007; Hu et al., 2018; McCormick et al., 2015; Pan et al., 2007; Steffen et al., 2008). This raises the question of whether the fall off in translation during normative aging is adaptive or maladaptive. Other studies implicate multiple facets of protein synthesis, including translational fidelity, ribosomal quality control, stoichiometric balance, aggregation, and turnover, as critical for cellular health (Anisimova et al., 2018; Dhondt et al., 2017; Koyuncu et al., 2021; López-Otín et al., 2023; Martinez-Miguel et al., 2021; Sacramento et al., 2020; Sirozh et al., 2024; Stein et al., 2022; Walther et al., 2015). Together, these observations suggest that selectively restoring ribosomal integrity later in life, rather than globally modulating translation, may benefit organismal homeostasis and longevity.

The nucleolus is a phase-separated organelle within the nucleus that coordinates ribosomal biogenesis (ribogenesis) through the orchestrated transcription and processing of ribosomal RNA (rRNA) and its assembly with RPs, and is therefore a major regulator of protein synthesis. We previously found that small nucleolar size is a cellular hallmark of longevity pathways across species, while enlarged nucleoli are associated with shorter lifespan. Notably, loss of the *C. elegans* cytosolic protein NCL-1, a homolog of BRAT/TRIM2/TRIM3, leads to enlarged nucleoli and disrupts longevity across diverse anti-aging paradigms, placing the nucleolus downstream of multiple life-extending pathways (Frank & Roth, 1998; Yi et al., 2015; Tiku et al., 2017). Consistent with this, human primary fibroblasts display a positive correlation between nucleolar dimension and chronological age, with patients suffering from Hutchinson-Gilford progeria syndrome, a rare accelerated aging disorder, exhibiting dramatically enlarged nucleoli (Buchwalter & Hetzer, 2017). Similarly, mammalian senescent cells as well as yeast mother cells near the end of their replicative life span display enlarged nucleoli, supporting nucleolar size as a biomarker of aging and senescence (Bemiller & Lee, 1978; Gutierrez & Tyler, 2024; Jo et al., 2024).

These findings raise the hypothesis that aging is associated with a progressive dysregulation of nucleolar function and ribosomal component production and assembly. In this framework, we used *ncl-1* loss-of-function mutants as a model of accelerated, age-related failure in ribosome biogenesis. We show that *ncl-1* loss disrupts ribosomal protein stoichiometry, decouples RP transcript and protein abundance, and compromises ribosomal subunit assembly, leading to loss of proteostasis. Importantly, we identify a means to robustly restore longevity downstream of nucleolar dysfunction by targeting either the RNAse P/MRP complex involved in rRNA and tRNA processing or the mitochondrial ribosome. These interventions sustain mature rRNA and RP abundance, improve cytosolic ribosomal subunit assembly, and reduce protein aggregation, despite persistent nucleolar enlargement and elevated pre-rRNA levels. Our findings demonstrate that restoring balance in ribosomal biogenesis, stoichiometry, and assembly can improve cytosolic ribosomal function and prolong lifespan.

## Results

### Loss of *ncl-1* in worms shortens lifespan and perturbs ribosome biogenesis

Loss of *ncl-1 (e1942)* expands nucleolar size and reduces the lifespan of major canonical longevity pathways (Tiku et al., 2017). To investigate the impact of enlarged nucleolar size specifically on somatic aging, we focused on *ncl-1 (e1942)* loss-of-function in long-lived *glp-1* (*e2141)* mutants, which contain somatic gonad but lack the germline. As previously shown (Tiku et al., 2017), *glp-1 ncl-1* double mutants exhibited significantly reduced longevity compared to *glp-1* single mutants, as well as increased senescence-associated β-galactosidase and reduced mobility at older ages (Figures 1A, 3F, S1C; Table S1), indicating that NCL-1 is essential for somatic health and longevity. Similarly, we found that *ncl-1* mutants on their own were shorter lived than wild-type N2 worms and had a steeper decline in age-dependent mobility, when exposed to the same temperature shifts as germlineless *glp-1* mutants, above (i.e. reared at 25°C and shifted to 20°C at day 1 of adulthood) (Figures 1B, S1A-B; Table S1), revealing that *ncl-1* mutation can cause premature aging in both *glp-1* and wild type contexts. This prompted us to ask whether nucleolar size changes during normative aging in wild-type N2 worms. Using fibrillarin (*fib-1::neongreen)* as a nucleolar marker, we observed a modest but significant increase in nucleolar size during adult aging (Figures 1C-D), corroborating similar findings in other species (Buchwalter & Hetzer, 2017; Gutierrez & Tyler, 2024).

**Figure 1.**
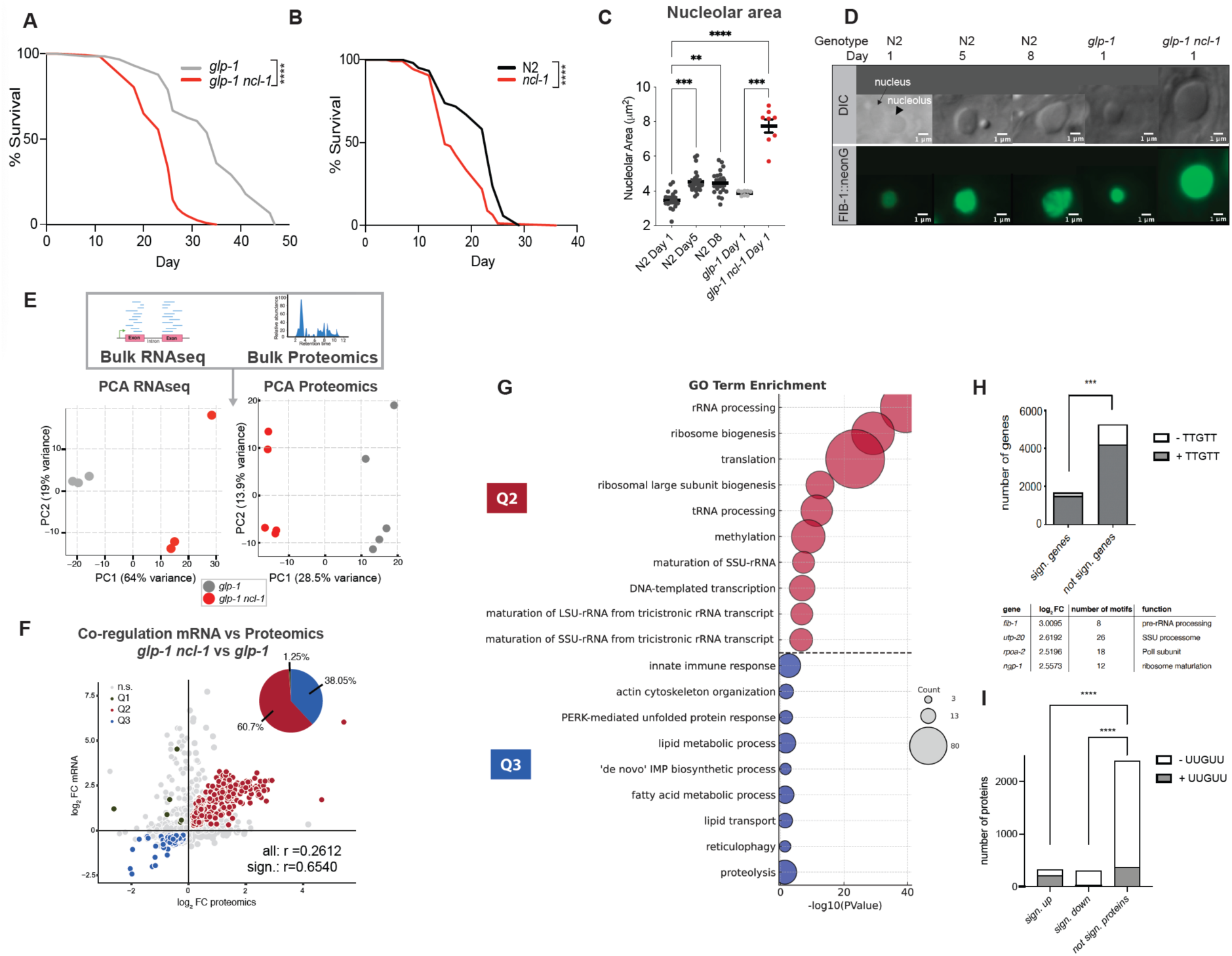
Disruption of *ncl-1* in germline-deficient *glp-1* worms reduces lifespan and elevates expression of genes involved in ribogenesis and protein synthesis A. Representative lifespan analysis of *glp-1 (e2141)* and *glp-1(e2141) ncl-1 (e1942)* worms. Lifespans were performed in ≥3 independent biological replicates, with n>100 worms per condition per experiment (Log-rank Mantel-Cox test, p<0.0001). B. Lifespan analysis of N2 and *ncl-1 (e1942)* worms. Lifespans were performed in ≥3 independent biological replicates, with n>100 worms per condition per experiment (Log-rank Mantel-Cox test, p<0.0001). C. Nucleolar area measurements of hypodermal cells in N2 (black) at day 1, 5, and 8 of adulthood, and *glp-1* (grey) and *glp-1 ncl-1* (red) at day 1 using FIB-1::neongreen to mark the nucleolus. Each dot represents the average of 5-15 nucleoli within one worm. (Kruskal-Wallis test post hoc Dunn’s multiple comparisons test, n =8-20 worms). N2 at day 5 vs N2 at day 1 p = 0.0003, N2 day 8 vs N2 at day 1 p = 0.0033, *glp-1 ncl-1* vs *glp-1* p = 0.0001, *glp-1 ncl-1* vs N2 =<0.0001, *glp-1* vs N2 n.s. D. Representative images of nucleoli by differential interference contrast microscopy (DIC) and FIB-1::neongreen fluorescence in head hypodermal cells. E. Principal-component analysis (PCA) of RNA-seq and LC-MS/MS data from *glp-1* (grey) and *glp-1 ncl-1* (red) at day 1. Each point corresponds to one biological replicate (RNA-seq n=3, proteomics n=5). The analysis was carried out on normalized gene-expression counts, and the first and second principal components (PC1, PC2) are shown. The clustering indicates that the two genotypes separate primarily along PC1. F. Co-regulation plot of transcript and protein logfold changes of *glp-1 ncl-1* vs *glp-1*. Quadrants are defined as Q1 post-transcriptionally repressed, Q2 co-upregulated, Q3 co-downregulated, and Q4 post-transcriptionally increased co-downregulated. Pie chart showing the percentage of candidate distribution in each quadrant, revealing most changes as co-regulated and a prevalence of up-regulated candidates. Colored candidates are significantly changed in both datasets with a p. adj<0.05. G. Top 10 GO: BP enriched in quadrant Q2 and Q3, derived from significant hits from E. (p. adj. <0.05). H. The NCL-1 mRNA-binding motif (TTGTT) is significantly enriched in the fraction of significantly regulated genes in *glp-1 ncl-1* mutants (Fisher’s exact test). Shown are representative genes with high fold change on the mRNA level and high motif number. I. Proteomic analysis reveals a strong, direction-specific enrichment of the NCL-1–associated post-stop motif (UUGUU) among proteins upregulated in *glp-1 ncl-1* mutants compared to *glp-1*. Stacked bar plots show the number of proteins with (+UUGUU, grey) or without (−UUGUU, white) the motif in significantly upregulated, significantly downregulated, and not significantly regulated protein groups. (Sign. upregulated vs not significant p = 3.74 x10^-70^, sign. downregulated vs not significant p = 1.66 x 10^-5^, Fisher’s exact test). ***<0.001, ****<0.0001

To uncover molecular mechanisms underlying *ncl-1* age acceleration, we performed bulk transcriptomic and proteomic analyses comparing *glp-1 ncl-1* double mutants to *glp-1* controls at maturity (day 1 adults). Differential gene expression analysis revealed widespread remodeling at both levels, showing 1633 transcripts significantly upregulated and 960 downregulated, alongside 331 upregulated and 320 downregulated proteins (adj. p-value < 0.05; Figures 1E, S1D-E; Table S2), with the overall correlation between the transcriptome and proteome comparable between genotypes (R² = 0.48 vs. 0.49; Figures S1F-G). Most significant changes in mRNA-protein correlation were either co-up or co-downregulated (R^2^ = 0.654, 98.1% in Quadrants 2/3 combined), with a clear prevalence of upregulation in the absence of *ncl-1*, consistent with the idea that NCL-1 post-transcriptionally represses mRNA and/or protein levels through binding to UUGUU motifs located in the 3’UTR (Figure 1F-G) (Sonoda & Wharton, 2001; West et al., 2018; Yi et al., 2015). Accordingly, this motif was significantly enriched (p<0.001, Fisher’s exact test) in the fraction of differentially regulated transcripts (Figures 1H, S1H). Among the genes with a high fold change and motif number were several factors involved in rRNA transcription, rRNA processing, and ribosome biogenesis, including *fib-1, utp-20, rpoa-2,* and *ngp*-1 (Figures 1H, S1I).

This motif enrichment extended to the proteomic level. Among significantly upregulated proteins in *glp-1 ncl-1* mutants, 62.5% contained three or more post-stop UUGUU motifs, compared to 15.3% among non-significantly regulated proteins (Figure 1I). In contrast, significantly downregulated proteins were depleted for the motif, with only 6.64% containing three or more UUGUU motifs (Figure 1I), revealing a strong direction-specific association between motif presence and increased protein abundance in the absence of *ncl-1*. Consistent with this pattern, enrichment analysis of proteins containing more than three UUGUU motifs demonstrated a significant bias toward upregulation in *glp-1 ncl-1* mutants (Figure S1J). Functional annotation of these upregulated, motif-rich proteins revealed strong enrichment for pathways related to ribosome biogenesis, rRNA and tRNA processing, and both cytosolic and mitochondrial translation (Figure S1K), indicating that loss of *ncl-1* preferentially enhances the abundance of translation-related proteins bearing its cognate regulatory motif.

More broadly, gene ontology analysis of the significantly altered genes in both RNA-seq and proteomics further illustrated the focal impact of *ncl-1* disruption on nucleolar related functions (Figure 1G; Table S3): Among the top 10 co-upregulated categories (Q2), we saw an enrichment of rRNA processing, ribosome biogenesis, translation (both cytosolic and mitochondrial), and tRNA aminoacylation. Co-downregulated categories (Q3) were enriched for lipid transport, fatty acid metabolic process, innate immune response, PERK-mediated stress response, and proteolysis. These data suggest that NCL-1 is a gatekeeper for ribosome biogenesis whose loss drives an accelerated aging state characterized by enlarged nucleoli, dysregulated ribosome biogenesis, and reduced somatic healthspan.

### Ribosomal proteins exhibit transcript-protein decoupling during aging

Progressive discoordination between mRNA and protein levels has been associated with loss of stoichiometric balance within protein complexes during aging (Di Fraia et al., 2025; Sacramento et al., 2020). To assess how aging impacts transcript-protein coordination in WT *C. elegans*, we integrated published single-worm RNA-seq data (Eder et al., 2024) with our own single-worm proteomics dataset from N2 animals sampled from day 1 to day 16 of adulthood (Figure 2A; Table S4). Weighted gene co-expression network analysis (WGCNA) was performed to identify gene modules exhibiting similar expression patterns and trajectories across aging within individual datasets, thereby capturing age-associated expression changes. Notably, among genes showing anticorrelated behavior, where transcript levels increased, but respective protein levels decreased with age, there was a significant enrichment of gene ontology terms associated with ‘translation’, ‘nucleolus’, and ‘ribosome’ (Figures 2B-C; Table S5). When specifically focusing on ribosomal proteins (RPs), we could see a significant negative correlation within a subset of the RP genes (Figures 2D-E), revealing a pronounced age-associated decoupling between ribosomal protein transcripts and their corresponding protein abundances. This pattern, previously observed in other organisms (Di Fraia et al., 2025; Khatir et al., 2023), represents the first such evidence of progressive RP decoupling during worm aging.

**Figure 2.**
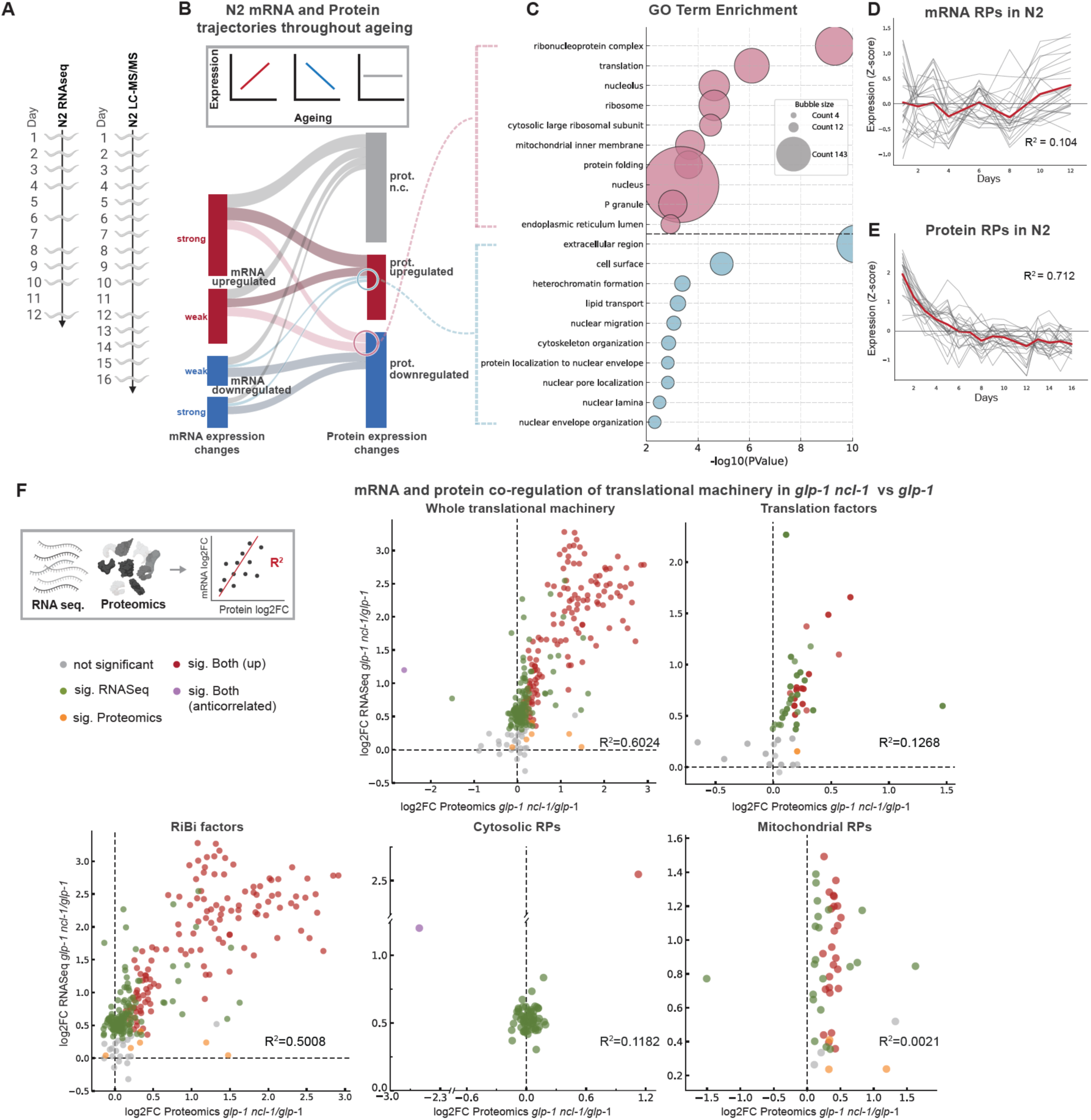
Age-associated decoupling of ribosomal protein mRNA and protein abundance **A.** Schematic illustrating experimental sampling for single-worm RNA sequencing (BioProject: PRJNA1015633, (Eder et al., 2024a)) and single-worm proteomics across adult aging. **B.** Schematic of mRNA–protein expression trajectories during ageing in wild-type N2 worms. Genes were clustered based on WGCNA modules (see Methods) as upregulated, downregulated, or not changed across adult ageing at the mRNA level (left) and the protein level (right). Bar colours represent direction of change (blue: downregulated; red: upregulated; grey: unchanged). The connecting flows indicate how each gene transitions between mRNA and protein categories. Color of each flow denotes the relationship between the two datasets: dark blue = downregulated at both mRNA and protein levels; dark red = upregulated at both levels; light pink = upregulated at mRNA but downregulated at protein level; light blue = downregulated at mRNA but upregulated at protein level. This representation highlights concordant and discordant ageing trajectories between transcript and protein abundance. **C.** Top 10 GO: BP and GO: CC enriched in the significantly anti-correlated mRNA-protein sets (light pink = upregulated at mRNA but downregulated at protein level; light blue = downregulated at mRNA but upregulated at protein level) derived from B. (p. adj. <0.05). Z-score expression trajectories of RP genes across ageing in wild-type N2 worms at the mRNA level, **D**. and **E**. protein level. Each line represents an individual RP; the red line shows the average trend. **F**. Co-regulation of transcript and protein logfold changes of the translational machinery categorized by GO: BP using data from Figure 1E-G. Grey represents genes that are not significant in both datasets, green are significant only in RNA-seq, orange only in proteomics, red are significant in the same direction in both datasets, pink are significant in both datasets but in opposite direction. (p values for correlations: whole translation machinery p <0.0001, cytosolic RPs p =0.0047, mitochondrial RPs p=0.7456, translation factors p <0.0005, and ribosome biogenesis p <0.0001.) ***<0.001, ****<0.0001

Building on this finding, we next examined whether this signature was altered by comparing *glp-1 ncl-1* and *glp-1* worms at day 1 of adulthood (Figure 2F). We first delineated the set of genes under defined GO biological processes, and assessed transcript-protein co-regulation for those detected at both mRNA and protein levels in our datasets. Ribosomal biogenesis factors involved in ribosome assembly and rRNA processing, as well as translation factors (initiation, elongation, and termination factors) generally showed a pronounced increase at both transcript and protein levels induced by *ncl-1* loss (R^2^ = 0.5008 and R^2^ =0.1268, respectively). In contrast, most cytosolic ribosomal proteins (RPs) showed transcriptional upregulation, which was not reflected at the proteome level, suggesting a loss of coordination between mRNA and protein for RPs (R^2^= 0.1182). (Exceptionally,

RPL-24.2 was significantly increased at both mRNA and protein levels, while RPL-11.1 mRNA levels were increased, but its protein abundance decreased). Mitochondrial RPs similarly exhibited minimal correlation between transcript and protein levels (R² = 0.0021). Notably, however, 48% of mitochondrial RPs were significantly upregulated in both datasets, and the lack of correlation appears to stem from the broad spread in transcript-level changes. Thus, while ribosome biogenesis and translation factors show coordinated transcriptional and proteomic upregulation in *glp-1 ncl-1* mutants, both cytosolic and mitochondrial RPs exhibit marked transcript-protein decoupling. The premature emergence of this aging-associated signature at day 1 supports the idea that *ncl-1* loss drives an accelerated aging state of ribosomal protein dysregulation.

### RNAi screen identifies mitochondrial ribosome and RNAse P/MRP complex components as suppressors of *ncl-1* aging phenotypes

To identify molecular pathways that mediate the accelerated aging phenotype in *ncl-1* loss-of-function mutants, we conducted a targeted RNAi suppressor screen. Based on our transcriptomic and proteomic analyses, we selected candidates falling under enriched GO terms that exhibited significant expression changes in *glp-1 ncl-1* mutants compared to *glp-1* controls. After additional filtering for conservation, UUGUU motif enrichment, and non-lethality, we focused on 67 candidates for functional validation (Figure 3A).

**Figure 3.**
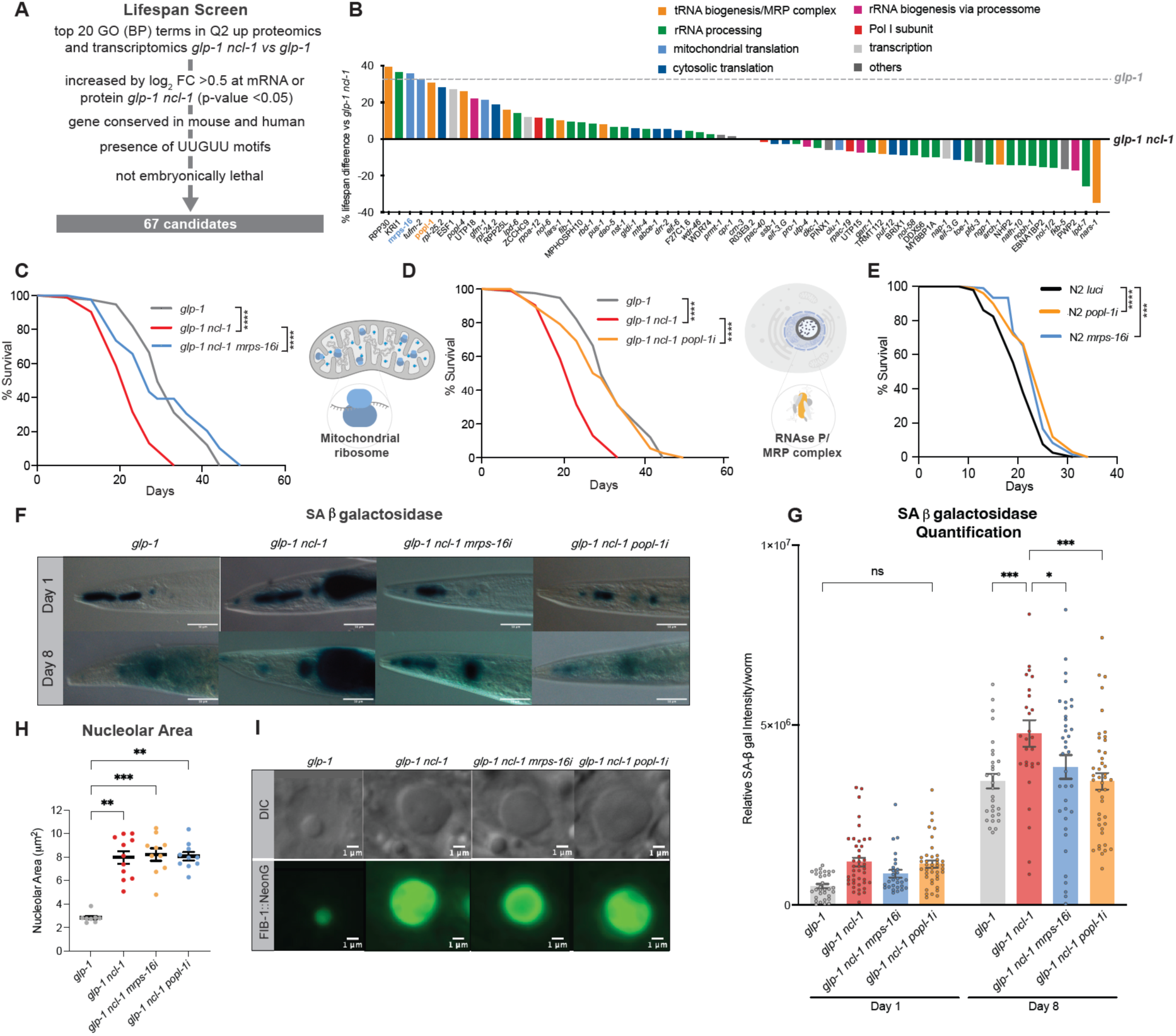
RNAi screen identifies mitochondrial ribosome and RNAse P/MRP complex components as suppressors of *ncl-1* aging phenotypes **A.** Candidate selection for RNAi lifespan rescue screen based on GO-term enrichment and fold changes at transcript and protein level in *glp-1 ncl-1* vs *glp-1*. Further filtering by the presence of UUGUU, conservation, RNAi availability, and lethality resulted in 67 candidates. **B.** Percentage lifespan difference of the RNAi treatments in *glp-1 ncl-1* relative to *glp-1 ncl-1* on luciferase control RNAi (set as 0). Grey dashed line representing the percent lifespan extension of *glp-1* compared to *glp-1 ncl-1*. RNAi treatments are color-coded depending on their respective GO-term (lifespans performed with n>100 worms). **C.** Lifespan analysis of *glp-1 ncl-1* worms on *mrps-16* RNAi, **C**. or *popl-1* RNAi **D.,** and control RNAi. Lifespans were performed in 5 independent biological replicates, with n>100 worms per condition per experiment (log-rank Mantel-Cox test, p<0.0001). **D.** Lifespan analysis of N2 worms on *mrps-16* RNAi or *popl-1* RNAi and control luciferase RNAi. Lifespans were performed in 5 independent biological replicates, with n>100 worms per condition per experiment (log-rank Mantel-Cox test, p<0.0001 and p<0.0005). **E.** Representative images of heads stained with SA-β-galactosidase in *glp-1*, *glp-1 ncl-1* worms on luciferase control, *mrps-16* RNAi, or *popl-1* RNAi at day 1 and 8 of adulthood. (Scale bars 100 μm). **F.** Quantification of the SA-β-gal staining. Each dot represents quantification from one worm. (n=30-45 worms, 2-way ANOVA post hoc Tukey’s test: Day 1 n.s.; Day 8 *glp-1* vs *glp-1 ncl-1* p=0.0005, *glp-1 ncl-1* vs *glp-1 ncl-1 mrps-16i* p=0.0184, *glp-1 ncl-1* vs *glp-1 ncl-1 popl-1i* p=0.0001; Geometric mean with 95% CI) **G.** Nucleolar area measured on the hypodermis of *glp-1, glp-1 ncl-1, glp-1 ncl-1* on *mrps-16* RNAi, *glp-1 ncl-1* on *popl-1* RNAi at day 5 of adulthood with FIB-1::neongreen. Each dot represents the average of 5-15 nuceloli per worm. (n=8 worms, Kruskal-Wallis test post-hoc Dunn’s multiple comparisons test, mean ± SEM). *glp-1 ncl-1* vs *glp-1* p = 0.0018*, glp-1 ncl-1* on *mrps-16* RNAi vs *glp-1* p = 0.0010, *glp-1 ncl-1* on *popl-1* RNA vs *glp-1* p = 0.0027. *glp-1 ncl-1* on *mrps-16* RNAi vs *glp-1 ncl-1* n.s, and *glp-1 ncl-1* on *popl-1* RNA vs *glp-1 ncl-1* n.s **H.** Representative images of nucleoli by DIC microscopy and FIB-1::neongreen fluorescence in hypodermal cells of *glp-1, glp-1 ncl-1, glp-1 ncl-1* on *mrps-16* RNAi, *glp-1 ncl-1* on *popl-1* RNAi at day 5 of adulthood. ns = not significant, *<0.05, **<0.01, ***<0.001, ****<0.0001

We assessed the ability of each candidate RNAi to rescue the shortened lifespan of *glp-1 ncl-1* mutants as a rigorous test of functional restoration. Notably, a subset of candidates significantly extended the lifespan of *glp-1 ncl-1* worms consistently, rescuing longevity to levels comparable to those of *glp-1* controls (Figure 3B; Table S1). Among the most effective suppressors were genes encoding components of two distinct molecular complexes: the RNAse P/MRP complex and the mitochondrial ribosome. Knockdown of multiple components from each complex reproducibly extended the lifespan of *glp-1 ncl-1* worms (Figures 3B, S2A-B, Table S1).

We focused on *mrps-16* and *popl-1* as representative components of these complexes, due to their robust and reproducible lifespan rescue effects in subsequent lifespan replicates (Figures 3C-D; Table S1). *mrps-16* encodes a small subunit protein of the mitochondrial ribosome, while *popl-1* encodes a major subunit of the RNAse P/MRP complex involved in pre-rRNA processing as well as tRNA maturation. Importantly, knockdown of *mrps-16* or *popl-1* extended lifespan in wild-type worms (Figure 3E), showing that these genes can influence longevity independent of the germline. Both *mrps-16* and *popl-1* transcripts harbor 2-3 UUGUU motifs after the stop codon (Table S6), and *mrps-16::mKate* and *mKate::popl-1* endogenous reporters were upregulated in worms upon *glp-1 ncl-1* loss (Figures S2F-G), indicating that they are direct or indirect post-transcriptional targets of NCL-1. We also examined whether the effects of these knockdowns were synergistic. Simultaneous knockdown of *mrps-16* and *popl-1* did not further extend lifespan beyond that observed with either knockdown alone, and neither knockdown altered the expression level of the other (Figures S2C-E; Table S1). Altogether, these findings indicate that NCL-1 coordinately regulates *mrps-16* and *popl-1*, which act in a similar genetic pathway converging on a shared downstream mechanism.

We then sought to identify mechanisms by which these RNAi interventions extend lifespan and assessed their impact on age-related physiology. We first measured senescence-associated β-galactosidase (SA-β-gal) activity in ageing worms, as a measure of senescence-related phenotypes (Figures 3F-G) (González-Gualda et al., 2021; Venz et al., 2024; Nonninger et al., 2025). SA-β-gal staining increased with age across all genotypes, including the long-lived *glp-1* mutants. Loss of *ncl-1* further elevated SA-β-gal activity at both young and old time points, indicating exacerbated senescence-like phenotypes, while *mrps-16i* or *popl-1i* significantly reduced SA-β-gal staining at older age, suggesting that the suppressors ameliorate senescence-related changes.

We next wondered whether the lifespan-extending RNAi treatments impact nucleolar expansion, which is another feature of senescence (Bemiller & Lee, 1978; Jo et al., 2024). As expected, *glp-1 ncl-1* mutants exhibited significantly enlarged nucleoli compared to *glp-1* controls at both day 1 and day 5 of adulthood (Figures 3H-I). Surprisingly, neither *mrps-16i* nor *popl-1i* significantly affected nucleolar size in *glp-1 ncl-1* mutants. This finding indicates that nucleolar size and lifespan can be uncoupled through modulation of downstream pathways.

### Knockdown of mitochondrial RP or RNAse P/MRP complex sustains rRNA and ribosomal protein abundance in *ncl-1* mutants

Nucleolar size is thought to correlate with the transcription of rRNA, which can be approximated by measuring levels of pre-processed rRNA precursors containing transcribed spacers (i.e., ITS, ETS) (Ding et al., 2022). To investigate whether nucleolar enlargement in *glp-1 ncl-1* mutants reflects increased precursor rRNA production, we quantified precursor rRNA levels by RNA-seq and qRT–PCR. Pre-rRNA abundance was significantly elevated in *glp-1 ncl-1* worms compared to *glp-1* controls and remained high despite *mrps-16* or *popl-1* knockdown (Figures 4A-B, S3A-C), indicating that nucleolar enlargement in *ncl-1* mutants is strongly associated with increased pre-rRNA transcription, while not excluding additional contributions from downstream processing or turnover. Consistent with this interpretation, knockdown of the RNA polymerase I subunit *rpoa-2*, previously shown to reduce nucleolar size in wild-type animals (Eberhard et al., 2013), also suppressed nucleolar enlargement in *glp-1 ncl-1* mutants (Figure S3D).

**Figure 4.**
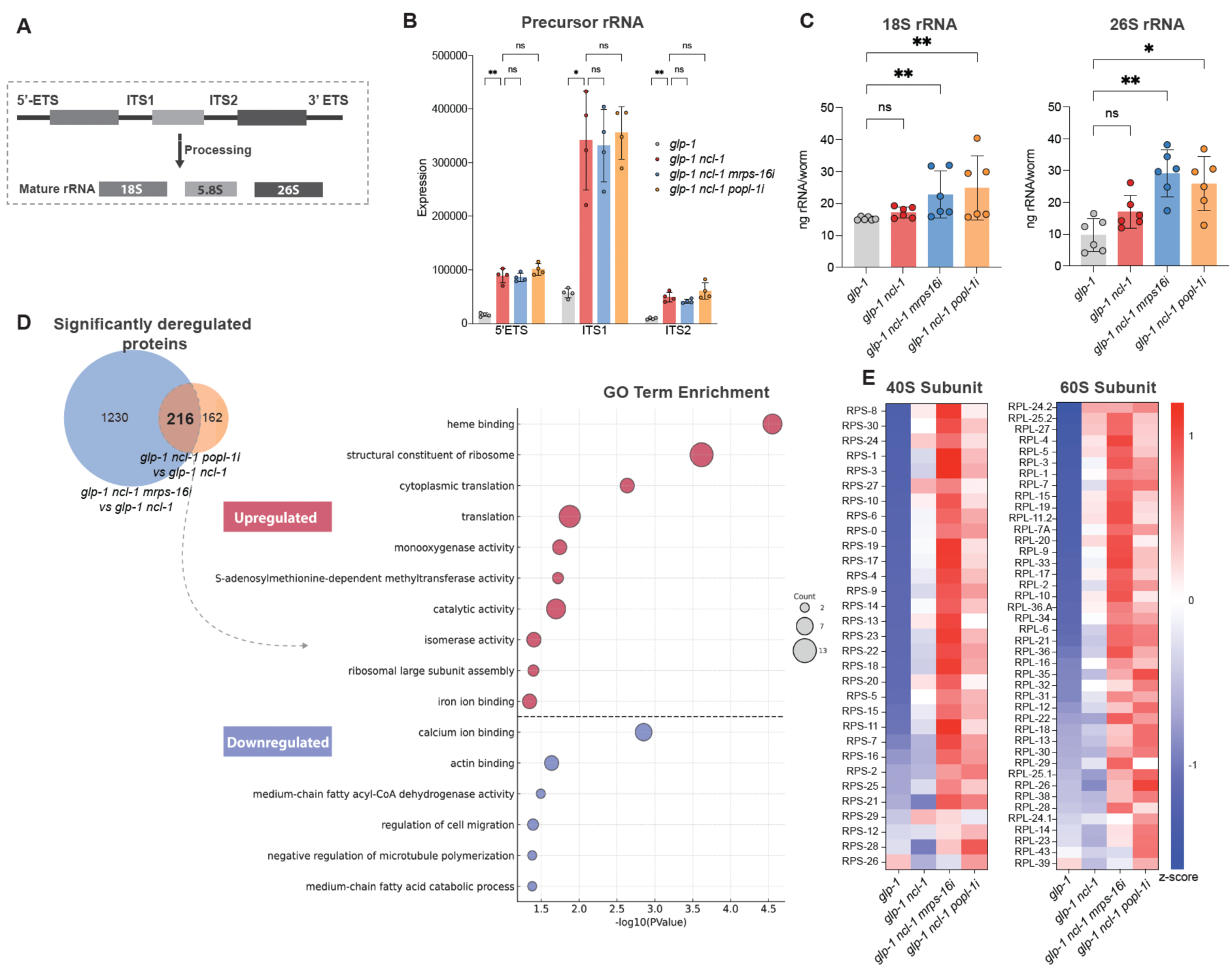
Knockdown of mitochondrial RP or RNAse P/MRP complex sustains rRNA and RP abundance. **A.** Schematic representation of the precursor rRNA containing ETS and ITS regions and its processing into mature 18S, 5.8S, and 26S rRNAs. **B.** Measurements of 5’ ETS, ITS1, and ITS2 from RNA-seq data in day 5 worms. *glp-1 ncl-1* worms have elevated levels of 5’ETS (p =0.0043), ITS1 (p =0.0285) and ITS2 (p =0.0069) compared to *glp-1*. *glp-1 ncl-1 on mrps-16i* or *popl-1i* do not have significantly different levels of 5’ETS, ITS1 or ITS2 compared to *glp-1 ncl-1*. Each dot represents a biological replicate. (n=4, 2-way ANOVA post hoc Tukey’s multiple comparisons test, mean ± SD). **C.** Mature 18S and 26S rRNA levels were measured at day 5. Mature rRNA levels of *glp-1 ncl-1* are not significantly different from *glp-1* on day 5*. glp-1 ncl-1* on *mrps-16i* has more 18S rRNA (p =0.0048) and 26S rRNA (p =0.0048) compared to *glp-1*. *glp-1 ncl-1* on *popl-1i* has significantly more 18S rRNA (p =0.0048) and 26S rRNA (p =0.0219) compared to *glp-1*. Each dot represents one independent experiment, with all genotypes measured within the same run. (n=6, Friedman ANOVA post hoc Dunn’s multiple comparisons test, mean ± SEM). **D.** Venn Diagram showing 216 common DEPs on day 5 of adulthood in *glp-1 ncl-1 mrps-16i* and *glp-1 ncl-1 popl-1i* compared to *glp-1 ncl-1*. Top enriched GO: BP terms from the overlapping significant DEPs (p adj. <0.05). **E.** Average Z-scores of all the cytosolic RPs detected in the single worm proteomics at day 5 in *glp-1*, *glp-1 ncl-1*, *glp-1 ncl-1 mrps-16i,* or *popl-1i*. For each protein, the mean across all samples was subtracted and divided by the standard deviation to derive the Z-score. 0 indicates expression equal to the protein’s average, positive values (red) higher-than-average, and negative (blue) lower-than-average. ns = not significant, *<0.05, **<0.01

Next, we assessed levels of mature rRNA by quantifying 18S and 26S rRNA species. At day 1 of adulthood, *glp-1 ncl-1* mutants displayed significantly increased 18S and 26S rRNA levels relative to *glp-1* controls, and treatment with *mrps-16i* or *popl-1i* had no further effect (Figures S4A-B).

However, by day 5, mature rRNA levels in *glp-1 ncl-1* worms dropped, and were, surprisingly, no longer significantly elevated relative to *glp-1* controls (Figure 4C), despite persistently elevated pre-rRNA levels, suggesting an age-dependent uncoupling between pre-rRNA transcription and steady-state mature rRNA levels. In contrast, *glp-1 ncl-1* worms treated with *mrps-16i* or *popl-1*i maintained significantly elevated 18S and 26S rRNA levels compared to controls, suggesting better coordination between rRNA production and maturation with age.

The preservation of mature rRNA levels in *glp-1 ncl-1* worms treated with *mrps-16i* or *popl-1i* raised the question of whether restored rRNA homeostasis reflects improved ribosome composition and proteome integrity. To address this, we performed single-worm proteomic analyses of *glp-1 ncl-1* mutants and RNAi-treated animals at day 1 and day 5 of adulthood (Figures S4C–F; Tables S7-S8). At day 5, differential expression analysis revealed extensive proteome remodeling in *glp-1 ncl-1* mutants compared to *glp-1* controls, with 1008 proteins significantly upregulated and 1036 downregulated (Figures S5A-D). Both *mrps-16i* and *popl-1i* treatments substantially altered the proteome of *glp-1 ncl-1* mutants, with 1446 and 378 differentially expressed proteins compared to *glp-1 ncl-1*, respectively. Comparing the proteome changes induced by the two suppressors indicated a significant overlap of 216 differentially expressed proteins (Figure 4D; Table S9), suggesting some common molecular pathways mediate their lifespan-extending effects. Gene ontology analysis of these shared targets revealed that commonly downregulated proteins were enriched for cytoskeletal organization, cell migration, and fatty acid metabolism. Interestingly, within the commonly upregulated proteins, there was a significant enrichment in processes related to ribogenesis, cytosolic and mitochondrial translation, and RNA processing (Figure 4D, Table S10). This finding indicates that both suppressors converge on similar processes, including regulation of the protein synthesis machinery, despite their residing in distinct complexes and compartments (mitochondria and nucleolus).

Given that gene ontology analysis of the shared proteomic changes induced by mrps-16i and popl-1i revealed enrichment for ribogenesis and translation-related processes (Figure 4D), we next examined whether ribosomal protein abundance itself is altered with age. Z-score visualization revealed that at day 1 of adulthood, *glp-1 ncl-1* mutants exhibited moderately increased abundance of multiple ribosomal proteins compared to *glp-1* controls (Figures S4G-H). However, this difference became less pronounced by day 5, where *glp-1 ncl-1* mutants showed reduced elevation, while *mrps-16i* and *popl-1i* showed sustained elevation of ribosomal proteins (Figure 4E), indicating maintenance of cytosolic ribosomal protein production. On the other hand, *popl-1i* had minimal effects on mitochondrial RP abundance, while *mrps-16i* selectively altered mitochondrial ribosomal subunit composition, consistent with its direct role (Figures S4I-J, S5E-F). Collectively, these results indicate that *popl-1i* and *mrps-16i* sustain higher levels of both cytosolic ribosomal proteins and mature rRNA with age, potentially supporting ribosomal function despite persistent nucleolar enlargement.

### Mitochondrial ribosome and RNAse P/MRP complex knockdown restores ribosomal protein stoichiometry and mRNA-protein coordination

We next asked whether, in addition to simply increasing ribosomal protein levels, the lifespan-extending suppressors actually preserve ribosomal stoichiometry and coordination during aging. To quantify the coordination among ribosomal proteins, we computed the pairwise protein–protein correlations among RPs from our proteomic data on day 1 and day 5 of adulthood (Figure 5A). As a negative control, we recalculated pairwise correlations using an abundance-matched background protein set (Leote et al., 2024) (Figures 5B-C). This analysis was performed across independent single-worm proteomics experiments, yielding consistent RP correlation distributions across batches (Figure S8A–B). At day 5, *glp-1 ncl-1* shifted the RP distribution toward lower correlations relative to *glp-1* (p <0.0001), indicating a loss of stoichiometric coordination within the ribosome. Consistent with their lifespan effects, *mrps-16i* and *popl-1i* treatments had higher mean correlation densities within the RP pairs, rescuing ribosomal stoichiometry coordination (p <0.0001). Analysis at the level of individual ribosomal proteins revealed that a subset of RPs exhibited pronounced average correlation loss in *ncl-1* mutants, which was commonly rescued by both knockdowns (Figure S8C–D). Heatmaps depicting each pairwise correlation also showed less correlation in *ncl-1* mutants, which was improved by the suppressors (Figure 5D). These findings indicate that *ncl-1-driven* loss of ribosomal protein and RiBi stoichiometry is functionally reversible by perturbation of either the mitochondrial ribosome or the RNAse P/MRP complex.

**Figure 5.**
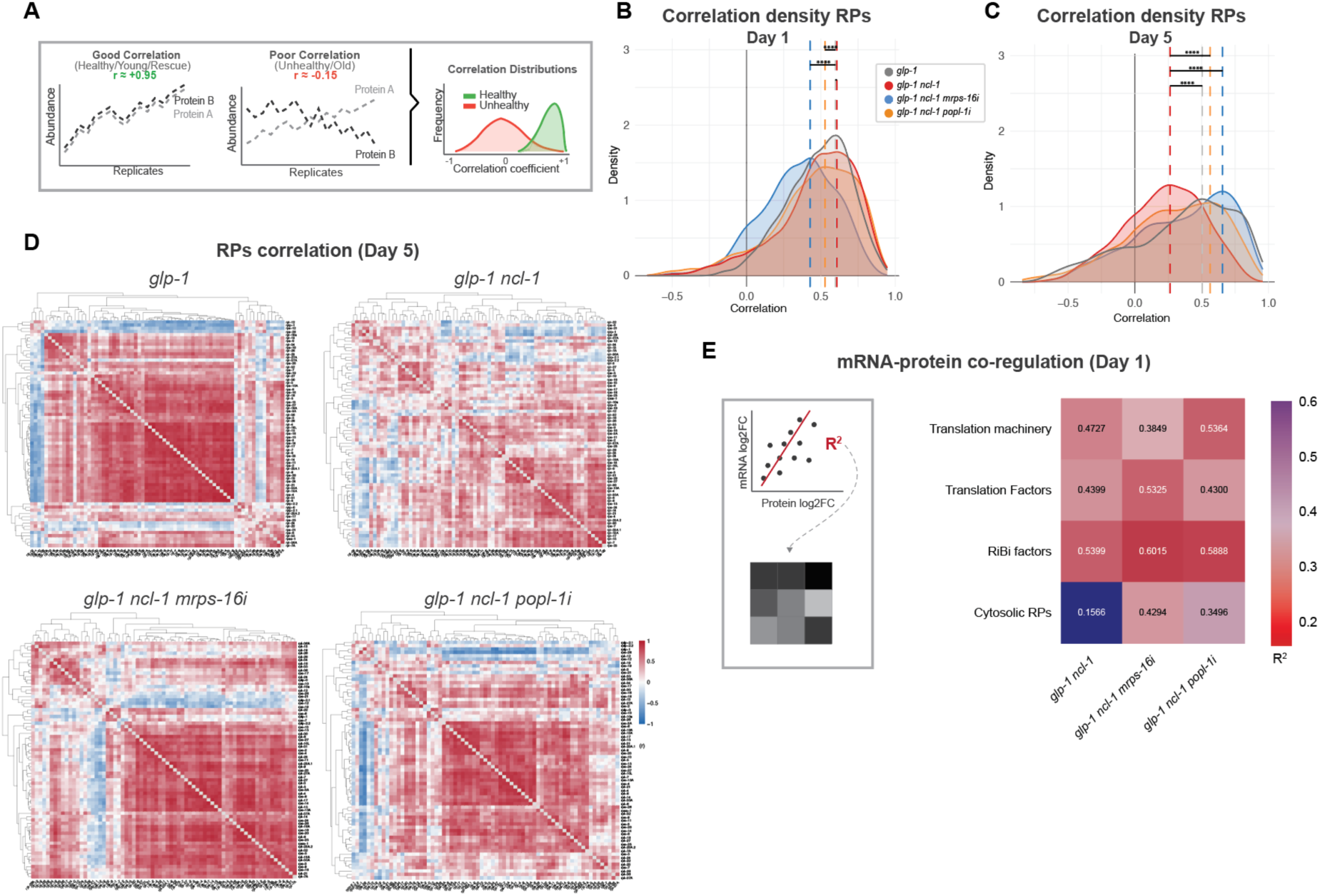
Mitochondrial ribosome and RNAse P/MRP complex knockdown restores ribosomal protein stoichiometry and mRNA-protein coordination. A. Protein abundance profiles with higher correlation (top) illustrate coordinated expression typical of healthy or youthful states, whereas lower correlation (bottom) reflects loss of coordination in unhealthy or aged states. The correlation coefficient distributions (right) highlight the global shift from high correlations in healthy samples to lower correlations in unhealthy samples. B. Distribution of pairwise protein-protein Pearson correlations among RPs at day 1 and **C**. day 5 in *glp-1*, *glp-1 ncl-1*, *glp-1 ncl-1 mrps-16i,* and *glp-1 ncl-1 popl-1i*. Dashed lines indicate the means of correlation for each condition. P-values denote two-sided Welch’s t-test comparing each condition to the corresponding glp-1 reference (All comparisons p<0.0001; ∼15 samples per experimental repeat, 3 experimental repeats were used for this analysis). C. Heatmap distribution showing the correlation (r) between each ribosomal protein in *glp-1*, *glp-1 ncl-1*, *glp-1 ncl-1 mrps-16i,* and *glp-1 ncl-1 popl-1i* at day 5. D. Co-regulation of transcript and protein (from single worm proteomics) R^2^ of the translational machinery categorized by GO: BP in day 1 worms. (Pearson R^2^ values; *glp-1 ncl-1* vs *glp-1*: whole translation machinery p <0.0001, cytosolic RPs p =0.0005 and mitochondrial RPs p=0.8054; *glp-1 ncl-1 mrps-16i* vs *glp-1 ncl-1*: whole translation machinery p <0.0001, cytosolic RPs p <0.0001 and mitochondrial RPs p=0.3973; *glp-1 ncl-1 popl-1i* vs *glp-1 ncl-1*: whole translation machinery p <0.0001, cytosolic RPs <0.0001 and mitochondrial RPs p=0.5814).

We next assessed whether this restored stoichiometry was also reflected at the level of mRNA–protein coordination. To do so, we analyzed transcript-protein co-regulation of the translational machinery using single-worm proteomic and transcriptomic data from day 1 of adulthood (Figures S6A-E), similar to Figure 2, above. Consistent with the bulk proteomics, *glp-1 ncl-1* mutants exhibited strong co-upregulation of the translational machinery as a whole (Figure 5E, top; R² = 0.473; Figure S6E). Within this grouping, ribosome biogenesis and translation factors showed the highest mRNA–protein concordance (Figure 5E, RiBi, R² = 0.54; translation factors, R² = 0.44; Figure S6E). In contrast, cytosolic and mitochondrial RPs displayed markedly weaker correlations (R² = 0.156 and R² = 0.0012, respectively), with ten cytosolic RPs showing anticorrelation. *mrps-16i* and *popl-1i* treatment induced comparable correlations of mRNA–protein coupling for translation and RiBi factors (Figures 5D, S6E; *mrps-16i* R² = 0.385 and 0.602; *popl-1i* R² = 0.536 and 0.589 for translation and RiBi, respectively). Importantly, *mrps-16i* and *popl-1i* knockdowns restored the correlation among cytosolic RPs (*mrps-16i* R² = 0.429; *popl-1i* R² = 0.350) and reduced the number of anticorrelated proteins to one (RPL-32) and two (RPL-12 and RPL-43). In contrast, mitochondrial RPs remained poorly correlated across all conditions, with *mrps-16* RNAi increasing the fraction of anticorrelated mitochondrial RPs (27.3% of the total number of mitochondrial RPs) relative to *glp-1 ncl-1* (5.4%) (Figure S6E). These results indicate that knockdown of either mitochondrial ribosome or RNAse P/MRP complex components restores cytosolic ribosomal protein stoichiometry and mRNA–protein coordination in *ncl-1* mutants. This restoration occurs despite persistent nucleolar enlargement, suggesting that improved ribosome composition and coordination are sufficient to extend lifespan in this accelerated-aging model.

### Loss of *ncl-1* disrupts ribosome assembly and proteostasis, which are restored by mitochondrial ribosome or RNAse P/MRP knockdown

To investigate whether the observed changes in ribosomal protein, rRNA levels, and ribosomal stoichiometries were associated with functional alterations in translation, we performed polysome profiling of the different strains at day 1 of adulthood (Figure 6A). Polysome profiles revealed accumulation of 40S and 60S ribosomal subunits in *glp-1 ncl-1* mutants compared to *glp-1* controls, accompanied by a reduction in 80S monosomes and polysomes, indicating a defect in ribosomal subunit joining (Figure 6B-C). Remarkably, both *mrps-16* and *popl-1* RNAi treatments normalized these profiles in *glp-1 ncl-1* worms, restoring the balance between free subunits and assembled ribosomes to levels comparable to those observed in *glp-1* controls. Thus*, ncl-1* mutants accumulate immature subunits that fail to assemble into functional ribosomes. These findings indicate that the impact of *ncl-1* loss impairs ribosomal subunit joining, a defect that is suppressed by downregulation of either the mitochondrial ribosome or the RNAse P/MRP complex.

**Figure 6.**
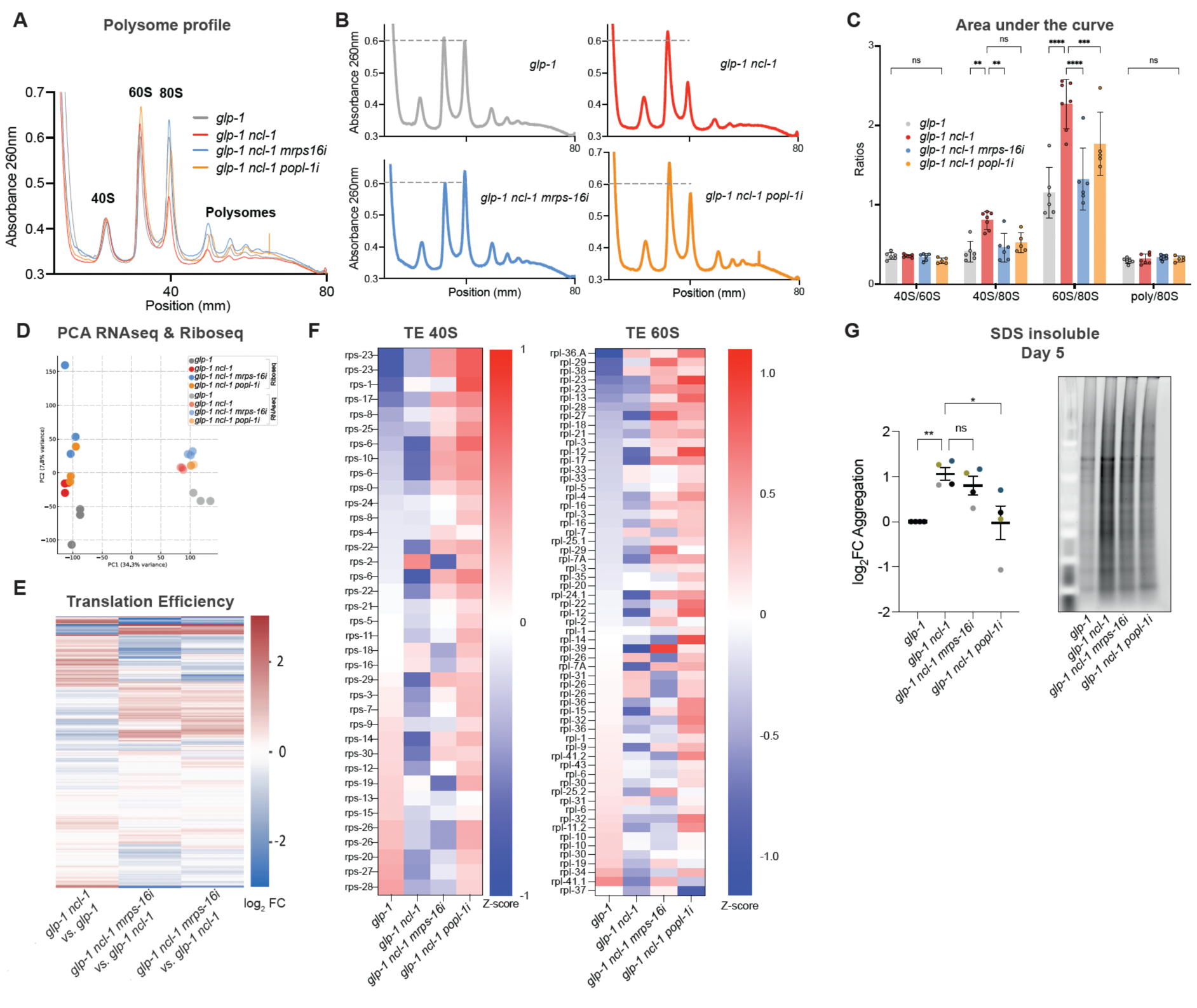
Loss of *ncl-1* disrupts ribosome assembly and proteostasis through translational misallocation, which is restored by mitochondrial ribosome or RNAse P/MRP knockdown. **A.** - **B**. Representative polysome profiles of day 1 *glp-1, glp-1 ncl-1, glp-1 ncl-1 on mrps-*16 RNAi and a*glp-1 ncl-1* on *popl-1* RNAi. Grey dashed line indicating the height of the 80S peak in *glp-1*. **B.** Quantification of the area under the curve from the polysome profiles in A. Each dot represents a biological replicate (n=6-7, 2way ANOVA post hoc Tukey’s test (40S/80S: *glp-1* vs *glp-1 ncl-1* p=0.0021, *glp-1 ncl-1* vs *glp-1 ncl-1 mrps-16i* p=0.0097, *glp-1 ncl-1* vs *glp-1 ncl-1 popl-1i* p=0.0683; 60S/80S: *glp-1* vs *glp-1 ncl-1* p<0.0001, *glp-1 ncl-1* vs *glp-1 ncl-1 mrps-16i* p<0.0001, *glp-1 ncl-1* vs *glp-1 ncl-1 popl-1i* p=0.0001) median with 95% CI). **C.** PCA of RNA-seq and Ribo-seq data from *glp-1* (grey), *glp-1 ncl-1* (red), *glp-1 ncl-1* on *mrps-16* RNAi (blue), and *glp-1 ncl-1* on *popl-1* RNAi (orange) at day 1. Each point corresponds to one biological replicate (n=3). The analysis was carried out on normalized gene-expression counts and ribo-counts, and the first and second principal components (PC1, PC2) are shown. The clustering indicates that the RNA-seq and Ribo-Seq data separate primarily along PC1, and the genotypes along PC2. **E.** Heatmap of the log2 fold-changes of the translation efficiencies of *glp-1 ncl-1* vs *glp-1, glp-1 ncl-1 mrps-16i* vs *glp-1 ncl-1,* and *glp-1 ncl-1 popl-1i* vs *glp-1 ncl-1,* separated by type of change categories. **F.** Average Z-scores of the translation efficiencies of all the cytosolic RP transcripts detected in the Ribo-seq in *glp-1*, *glp-1 ncl-1*, *glp-1 ncl-1 mrps-16,* or *popl-1* RNAi. For each transcript, the mean across all samples was subtracted and divided by the standard deviation. 0 indicates expression equal to the transcript’s average, positive values (red) higher-than-average, and negative (blue) lower-than-average. **G.** Quantification (left) and representative gel (right) of SDS-insoluble protein fractions from day 5 adult worms. *glp-1 ncl-1* mutants show increased aggregation relative to *glp-1* controls (p = 0.0078), which is reduced by *popl-1* RNAi (p = 0.0411). Dots represent individual biological replicates, colored by replicate identity (n = 4; one-way ANOVA post hoc Friedman’s test) ns = not significant, *<0.05, **<0.01, ***<0.001, ****<0.0001

Despite the altered polysome profiles, overall protein synthesis rates, assessed by the SUnSET assay, showed no statistically significant differences between the experimental groups at either day 1 or day 5 of adulthood after normalization per worm (Figures S7A-C). As a positive control, *rsks-1* mutants displayed the expected reduction in protein synthesis (Pan et al., 2007; Derisbourg et al., 2021). Additionally, worm size itself was unchanged across conditions (Figure S7D). Consistent with the idea of no difference in protein synthesis rates, the polysome-to-monosome ratio was unchanged across experimental groups. Together, these data indicate that although ribosomal subunit assembly is compromised in *glp-1 ncl-1* mutants, global translation rates are buffered. Such decoupling between ribosome biogenesis and bulk translation has been previously reported, including in ribosomal protein mutants with subunit-joining defects but preserved translational output (Kondrashov et al., 2011; Tiruneh et al., 2013) and upon disruption of specific RiBi factors that destabilize ribosome production without markedly reducing overall protein synthesis (Gómez-Herreros et al., 2017; X. Zhu et al., 2019).

Given preserved global translation, we next tested whether translational output is redistributed across specific mRNA classes. To this end, we performed ribosome profiling to measure translation efficiency (TE) in the *ncl-1* loss-of-function background and the *mrps-16* and *popl-1* RNAi rescues at day 1 of adulthood. Loss of *ncl-1* resulted in extensive translatome reprogramming, with 974 transcripts showing significant TE changes compared to *glp-1* controls (Figures 6D, S7E-I; Table S11). Among other things, ribosome biogenesis and rRNA processing transcripts showed increased TE, whereas ribosomal protein mRNAs were significantly downregulated at the translational level (Figure S7H; Tables S12-S13). This redistribution of translational output aligns with the proteomic changes we observed in *glp-1 ncl-1* double mutants (Figure 4D). In parallel, TE was reduced for transcripts associated with differentiated somatic functions, including signaling and ion transport, indicating a shift of translational resources away from tissue maintenance programs. Consistent with their rescue of the polysome phenotype, both *mrps-16* and *popl-1* RNAi largely reversed this translational misallocation. Compared to *glp-1 ncl-1* mutants, RNAi-treated animals showed restored TE of RP transcripts (Figure 6F) and increased TE of genes involved in proteasome-mediated protein catabolism, lipid metabolism, and organismal growth (Figures S7G, S7I). This shift aligns with the recovery of RP abundance and improved mRNA-protein coupling observed upon RNAi treatment (Figures 3E, 4D).

The increased TE for proteasome/ubiquitin-dependent catabolism suggests enhanced proteostasis capacity. We therefore next examined protein aggregation as a functional readout of proteostasis. We isolated SDS-insoluble proteins and quantified them in-gel (Figure 6G). Loss of *ncl-1* led to increased SDS-insoluble protein aggregates by day 5 of adulthood, whereas both mrps-16 and popl-1 RNAi reduced aggregate burden.

Together, these results indicate that loss of *ncl-1* impairs ribosomal subunit joining and proteotasis by misdirecting translation toward early ribosomal biogenesis at the expense of ribosomal protein production and cellular maintenance. Imbalanced stoichiometry exerts a constant proteostatic load that foments protein aggregation. Suppression of either mitochondrial ribosome or the RNAse P/MRP complex restores translational balance, ribosome assembly, and proteome integrity, providing a mechanistic explanation for their lifespan-extending effects in *ncl-1* mutants.

## Discussion

Ribosomal biogenesis, ribosomal protein abundance and global protein synthesis decline with age (Anisimova et al., 2020; Blazejowski & Webster, 1983; Brown et al., 2018; Dhondt et al., 2017; Di Fraia et al., 2025; Dwyer et al., 1980; Fando et al., 1980; Layman et al., 1976; Llewellyn et al., 2023; Lu et al., 2023; Motizuki & Tsurugi, 1992; Stein et al., 2022; Ubaida-Mohien et al., 2019; Walther et al., 2015; Ward & Richardson, 1991; Webster & Webster, 1979; H. Zhu et al., 2023), yet paradoxically, genetic or pharmacological downregulation of the translational machinery, particularly early in life, extends lifespan across species (Chen et al., 2007; Chiocchetti et al., 2007; Bitto et al., 2016; Curran & Ruvkun, 2007; Demontis & Perrimon, 2010; Hansen et al., 2007; Hu et al., 2018; McCormick et al., 2015; Pan et al., 2007; Steffen et al., 2008). Our findings reconcile these observations by showing that shortened lifespan is not caused by excessive ribosomal output, but instead by dysregulated ribosome biogenesis and impaired ribosome assembly. Using *ncl-1* loss-of-function mutants as a model of accelerated aging, we demonstrate that disrupted ribosomal protein stoichiometry, defective subunit joining, and translational misallocation characterize the short-lived phenotype. Our results, particularly in the longer-lived *glp-1* background, indicate that restoring ribosomal protein production and assembly can improve proteostasis and extend lifespan. This suggests that maintaining ribosomal integrity and stoichiometric balance, rather than broadly reducing overall protein synthesis, may be an important factor influencing longevity.

Our work reframes the relationship between nucleolar size, ribosome biogenesis, and longevity by placing *ncl-1* mutants within a broader framework of RiBi dysfunction observed in both aging and congenital disorders. While enlarged nucleoli and increased rRNA accumulation have often been interpreted as markers of enhanced rRNA production, ribosomal output, and protein synthesis, *ncl-1* mutants instead exhibit decoupling between nascent and mature rRNA, ribosomal protein transcripts, and protein levels, loss of RP stoichiometry, and inefficient subunit joining. These defects resemble age-associated changes reported across species (Anisimova et al., 2020; Di Fraia et al., 2025; Khatir et al., 2023; Sacramento et al., 2020) and also the cellular consequences of some ribosomopathies (e.g., Diamond–Blackfan anemia, Shwachman–Diamond syndrome, cartilage hair hypoplasia (Kang et al., 2021). Similarly, a progeroid mouse model with increased nucleolar stress showed an accumulation of free ribosomal proteins, impaired RiBi, and accelerated aging phenotypes, further linking nucleolar dysfunction and organismal decline (Sirozh et al., 2024).

Notably, despite these profound defects in RiBi, we do not detect significant changes in bulk protein synthesis. This suggests that global translation is buffered in *ncl-1* mutants and that the primary defect is qualitative rather than quantitative.

Importantly, we find that the shortened lifespan of *ncl-1* mutants can be rescued downstream of the nucleolus by rebalancing mature rRNA and ribosomal proteins and thus improving subunit assembly, proteostasis, firmly linking restored coordination to improved survival. These data support the view that qualitative failures in RiBi, specifically transcript–protein decoupling and stoichiometric imbalance, are central drivers of aging, rather than uniform declines in overall protein synthesis.

Studies in the short-lived killifish *N. furzeri* have shown that during aging, ribosome stalling increases and there is a progressive loss of coordination between transcript and protein abundance, especially for RPs, resulting in stoichiometric imbalance and impaired ribosome assembly that contribute to a widespread disruption of proteostasis (Di Fraia et al., 2025; Sacramento et al., 2020). We observe a similar uncoupling between RP transcripts and protein abundance in wild-type worms during aging, and already in young *ncl-1* mutants, consistent with their accelerated aging phenotype.

Aging is also associated with loss of stoichiometric balance within multiprotein complexes. In our analysis, ribosomal stoichiometry was assessed by quantifying the coordination among ribosomal proteins while controlling for abundance-driven effects using per-condition, abundance-matched background correlations. While this approach shows loss of stoichiometric balance within ribosomes in *ncl-1* mutants, it also suggests other complexes may be vulnerable to failure. At a broader systems level, proteomic analysis in humans has similarly reported age-associated declines in the coordinated regulation of protein complex stoichiometry across fundamental cellular processes (Leote et al., 2024). In agreement with our *ncl-1* data, proteomic profiling across lifespan in *C. elegans* has revealed a decline and imbalance, particularly in RPs and ribosomes (Walther et al., 2015). These changes are also accompanied by loss of proteostasis and increased aggregation, particularly of ribosomal proteins (Reis-Rodrigues et al., 2012).

Our results position NCL-1 as a central regulator of translational homeostasis in *C. elegans* aging. NCL-1 is a cytosolic RNA-binding protein with an established role in controlling nucleolar size through regulation of rRNA synthesis via FIB-1 (Frank & Roth, 1998; Tiku et al., 2017; Yi et al., 2015). NCL-1 belongs to the TRIM-NHL family of proteins, being closely related to Brat and Mei-p26 in *Drosophila* and TRIM2, 3, and 32 in mammals (Connacher & Goldstrohm, 2021). These proteins are known translational repressors acting via 3ʹUTRs (Kondrashov et al., 2011; Loedige et al., 2013, 2014, 2015; Salerno-Kochan et al., 2022). In worms, NCL-1 preferentially targets U-rich motifs in the 3’ UTR of transcripts encoding mainly proteins belonging to ribosomal biogenesis and the translation machinery (West et al., 2018). Accordingly, in our transcriptomic data, there is an enrichment for genes containing the TTGTT motif in the *ncl-1* mutants. Importantly, our proteomic analyses demonstrate that this motif-dependent regulation is propagated to the protein level, with UUGUU-containing targets showing a strong, direction-specific enrichment among upregulated proteins in *ncl-1* mutants, while being depleted among downregulated proteins. Notably, these targets include components of both cytosolic and mitochondrial ribosome biogenesis and translation, linking NCL-1-dependent post-transcriptional regulation to multiple ribosomal systems. Though communication between mitochondrial and cytosolic ribosome biogenesis has been noted previously (Molenaars et al., 2020; Suhm et al., 2018), our studies pinpoint NCL-1 as a central coordinator of these processes, highlighting their deep functional interrelationship.

Among NCL-1-regulated proteins, the mitochondrial ribosome (*mrps-16*) and RNAse P/MRP complex (*popl-1*) emerged because of their reproducible beneficial effects on the lifespan of *ncl-1* mutants and wild-type worms. A key feature of *mrps-16* and *popl-1* knockdown is that they restore ribosomal protein abundance without increasing global protein synthesis. Instead, they enhance the efficiency with which ribosomal protein mRNAs are translated. This mechanism explains how RP upregulation is achieved in the absence of increased ribosomal output, and places translational efficiency and allocation at the center of ribosome restoration and lifespan extension. These suppressors likely reflect processes downstream of the nucleolus that help coordinate the cellular translational machinery to match the cell’s metabolic and proteostatic state. Mechanistically, partial perturbation of mitochondrial translation and RNAse P/MRP may act through complementary routes converging on the cytosolic ribosome. Perturbation of mitochondrial translation is known to trigger mitonuclear protein imbalance and activate conserved longevity programs, such as the mitochondrial unfolded protein response, reprogramming stress signaling, and proteostasis that restore balance (Houtkooper et al., 2013; Liu et al., 2020; Soto et al., 2022). Mitochondrial stress signaling can also influence cytosolic translation through coordinated translational balancing mechanisms dependent on ATF-5/ATF-4 (Suhm et al., 2018; Molenaars et al., 2020). Perturbations in RNAse MRP and RNAse P, which set early pre-rRNA and tRNA processing kinetics, are known to reshape ribosome biogenesis, indicating that changes in processing rates can stabilize rRNA maturation and RP assembly (Chamberlain et al., 1996; Lindahl et al., 2009). The net effect is a more balanced coordination between rRNA maturation and RP supply, improved subunit export and joining, and reduced aggregation, consistent with restored proteostasis and extended lifespan. Whether *mrps-16* and *popl-1* extend lifespan in wild-type animals through the same ribosome biogenesis restoring mechanisms identified in *ncl-1* mutants remains to be determined. Notably, the beneficial effects in wild-type animals arise when knockdown is initiated in adulthood, suggesting that attenuation of these pathways becomes advantageous as age-associated decline emerges. This is consistent with *ncl-1* mutants, which exhibit accelerated aging and therefore require intervention at earlier stages.

Collectively, our data indicate that longevity depends on coordinated ribosome homeostasis rather than nucleolar size or ribosome abundance alone. This finding could explain why up- or downregulating RPs could have beneficial effects on life span dependent on context. Long-lived *glp-1* mutants maintain reduced but well-matched levels of ribosomes, ribosomal protein mRNAs and proteins, and mature rRNA, whereas *ncl-1* mutants display a dysregulated state with excess immature rRNA, disproportionate RP transcript accumulation, insufficient RP protein production, and defective assembly. Knockdown of the RNAse P/MRP complex or the mitochondrial ribosome restores this balance by generally upregulating RPs and increasing coordination, supporting proteostasis and lifespan extension without normalizing nucleolar size.

Together, prior work across species has established that nucleolar size inversely correlates with lifespan, and that typical ribosome aging is marked by loss of coordination and stoichiometry. Here, we demonstrate that these age-related defects can be reversed: in *ncl1* mutants, longevity can be restored without correcting nucleolar size. This refines the simple inverse correlation between nucleolar size and longevity (Buchwalter & Hetzer, 2017; Tiku et al., 2017), and suggests that interventions targeting ribosomal biogenesis and its assembly, rather than nucleolar size alone, may effectively mitigate age-related decline.

## Methods

### Experimental model and study participant details

#### C. elegans strains

The following strains were used: N2 (wild type), *glp-1(e2141)* III*, glp-1(e2141) ncl-1(e1942)* III*, ncl-1(e1942)* III*, fib-1(syb828)* V*, glp-1(e2141) ncl-1(e1942)* III*; fib-1(syb828)* V*, glp-1(e2141)* III *;fib-1(syb828) V, mrps-16(syb2433)* II*;glp-1(e2141)* III*, mrps-16(syb2433*) II*; glp-1(e2141) ncl-1 (e1942)* III*, popl-1(syb2735) glp-1(e2141)* III*, popl-1(syb2735) glp-1(e2141) ncl-1 (e1942)* III*, rsks-1(sv31)* III.

Previously published mutant strains were obtained from the Caenorhabditis Genetics Center (CGC) or the National BioResource Project (NBRP) and outcrossed to N2 at least twice before use.

Endogenously tagged strains were generated by CRISPR–Cas9 (Sunybiotech) and validated by Sanger sequencing.

#### Nematode culture conditions and maintenance

*C. elegans* strains were maintained on nematode growth medium (NGM) agar plates seeded with *Escherichia coli* OP50 at 20°C unless otherwise specified, following established methods (Brenner, 1974; Stiernagle, 2006). Strains carrying the temperature-sensitive *glp-1(e2141)* allele were maintained at 15°C. To induce sterility in *glp-1(e2141),* eggs were shifted to 25°C for 52 h until adulthood before use in experiments or returned to 20°C for lifespan analyses. Worms were transferred to fresh plates every few days to avoid starvation and contamination.

Synchronous populations were obtained either by timed egg-lay or by alkaline hypochlorite treatment. For timed egg-lay, gravid adults were placed on a fresh NGM plate seeded with the corresponding bacteria strain and allowed to lay eggs for 3–4 h at the experimental temperature; adults were then removed. For bleaching, gravid adults were washed from plates with M9 buffer, pelleted by low-speed centrifugation, and treated with freshly prepared alkaline hypochlorite solution until adult carcasses were lysed while eggs remained intact. Eggs were recovered by centrifugation, washed twice with M9 buffer, and plated onto NGM plates seeded with OP50.

## Method details

### Lifespan assay

Worms used for lifespan analysis were synchronized by egg laying. Assays were performed at 20 C or 25 C to induce the *glp-1(e2141)* genotype. At least 100 worms per strain were used, with 20-30 worms per 6 cm plate with the corresponding bacteria RNAi strain. Worms were transferred to fresh plates every three days or every day. Scoring of deaths and censoring (e.g., because of internal hatching, vulval protrusion, or burrowing in agar) was carried out every second or third day. Mean, median, and maximum lifespan were calculated and plotted using Prism GraphPad.

### RNA interference

RNAi by feeding was performed using standard protocols (Stiernagle, 2006). *E. coli* HT115(DE3) bacteria carrying double-stranded RNA expression plasmids were obtained from the Vidal (Rual et al., 2004) or Ahringer (Kamath & Ahringer, 2003) RNAi libraries. HT115(DE3) bacteria carrying a luciferase RNAi construct (L4440) were used as a control. RNAi bacteria were grown overnight in lysogeny broth containing 50 µg/mL ampicillin and seeded onto RNAi plates supplemented with 1 mM isopropyl β-D-1-thiogalactopyranoside (IPTG) and 50 µg/mL ampicillin. RNAi was initiated by transferring synchronized animals onto RNAi plates, typically from the egg stage unless otherwise specified.

### Mature rRNA measurements

Total RNA was extracted from synchronized adult worms using a QIAzol–RNeasy–based protocol. For each condition, 100 worms were collected in M9 buffer, flash-frozen in liquid nitrogen, and stored at −80°C until processing; 400 N2 worms were processed in parallel to generate a standard curve. Samples were homogenized by repeated freeze–thaw cycles in QIAzol, followed by chloroform phase separation and purification on RNeasy Mini spin columns (Qiagen) according to the manufacturer’s instructions.

RNA concentration of N2 samples was determined by NanoDrop and used to generate a standard curve. Equal volumes of RNA from experimental samples were resolved alongside defined amounts of N2 RNA (0.125–1 µg) on 1% agarose gels stained with RotiRed. Mature rRNA bands were visualized and quantified by densitometry, and rRNA abundance in experimental samples was interpolated from the standard curve.

### Quantitative RT-PCR

Synchronized adult worms were collected into ice-cold M9 buffer, pelleted, flash-frozen in liquid nitrogen, and lysed by repeated freeze–thaw cycles. Lysates were clarified by centrifugation, and RNA was extracted using QIAzol followed by purification on RNeasy Mini spin columns (Qiagen) according to the manufacturer’s instructions. RNA concentration and purity were assessed by NanoDrop spectrophotometry.

cDNA was synthesized from total RNA using the iScript cDNA Synthesis Kit (Bio-Rad). Quantitative PCR was performed using Power SYBR Green Master Mix (Applied Biosystems) on a ViiA 7 Real-Time PCR System. Relative transcript levels were calculated using the ΔΔCt method, with *snb-1* or *etf-3* used as endogenous controls.

### RNA-Sequencing

Total RNA was extracted from synchronized adult worms and purified as described above. Libraries were prepared from 300 ng total RNA. Enzymatic depletion of ribosomal RNA with the Illumina Ribo-Zero Plus rRNA Depletion Kit was followed by library preparation with the Illumina® Stranded Total RNA sample preparation kit. The depleted RNA was fragmented and reverse transcribed with random hexamer primers, second-strand synthesis with dNTPs was followed by A-tailing, adapter ligation,n and library amplification (12 cycles). Next, the library validation and quantification (Agilent Tape Station) were performed, followed by pooling of equimolar amounts of the library. The pool itself was then quantified using the Peqlab KAPA Library Quantification Kit and the Applied Biosystems 7900HT Sequence Detection System and sequenced on an Illumina NovaSeq6000 sequencing instrument with a PE100 protocol aiming for 25 million clusters per sample.

Reads were mapped to the *Caenorhabditis elegans* reference genome (Ensembl 110) using Kallisto (v0.46.1). After normalization of read counts by making use of the standard median-ratio for estimation of size factors, pair-wise differential gene expression analysis was performed using DESeq2 (v1.24.0). After removal of genes with fewer than 10 overall reads, logfold changes were shrunken using approximate posterior estimation for GLM coefficients.

Principal component analysis and enrichment visualization were generated using Flaski (v2.3.0), an interactive web application suite for biological data analysis provided by the Bioinformatics Core Facility at the Max Planck Institute for Biology of Ageing (Iqbal et al., 2021). P values from Fisher’s exact tests were adjusted for multiple hypothesis testing; adjusted P < 0.05 was considered significant.

### NCL-binding motif analysis

Enrichment of the NCL-1 RNA-binding motif (UUGUU) among genes regulated in *glp-1 ncl-1* mutants was assessed using custom Python scripts. Transcript sequences were extracted from annotated transcript models using bedtools, and UUGUU motif occurrences were identified using regular expression–based searches. Motif counts were summarized per gene and merged with RNA-seq differential expression results comparing *glp-1(e2141)* and *glp-1(e2141) ncl-1(e1942)* animals.

Enrichment was evaluated by comparing motif abundance in significantly versus non-significantly regulated genes using Fisher’s exact test.

### Bioinformatic precursor rRNA analysis

To quantify precursor and mature ribosomal RNA (rRNA) from the RNA-seq data, we constructed an augmented reference that includes the full ribosomal DNA (rDNA) repeat unit. We obtained the complete *C. elegans* 45S rDNA sequence (including the 18S, 5.8S, and 26S rRNA coding regions along with spacer sequences) and associated annotations from NCBI (accession MN519140.1). Because rDNA repeats are not fully represented in standard genome assemblies, we masked the endogenous tandem rDNA cluster (Ensembl 110 coordinates: chr I: 15,057,500–15,072,434) in the reference before augmentation. The rDNA sequence and its annotation were concatenated with the masked WBcel235 reference FASTA and GTF to produce a combined reference suitable for read mapping.

Trimmed RNA-seq reads were aligned to the augmented reference using STAR with settings optimized for multimapping across repetitive sequences. Counts were assigned to rDNA features (18S, 5.8S, 26S, and spacer regions) using the feature. Because the augmented reference includes the complete rDNA unit, read counts mapping to these coordinates capture both precursor and mature rRNA transcripts derived from the rDNA loci (Ellis et al., 1986).

Count matrices for rRNA features were then integrated with the gene expression dataset and analyzed using DESeq2 for normalization and differential expression between day 1 and day 5 samples. Normalized rRNA read counts were used to assess relative changes in rRNA abundance, with statistical significance determined using DESeq2’s standard multiple testing correction (adjusted P < 0.05).

### Bulk proteomic sample preparation and analysis

Age-synchronized day 1 adult worms were collected from NGM plates (six plates per condition), washed extensively to remove bacteria, and lysed in LC–MS/MS–compatible buffer by heat denaturation and Bioruptor sonication. Lysates were clarified by centrifugation, and protein concentration was determined. For each sample, 300 µg total protein was digested with trypsin (1:200, enzyme: protein). Peptides were desalted on C18 StageTips and dried before LC–MS/MS analysis.

Peptides were separated on a 25 cm, 75 μm internal diameter PicoFrit analytical column (New Objective) packed with 1.9 μm ReproSil-Pur 120 C18-AQ media (Dr. Maisch) using an EASY-nLC 1200 (Thermo Fisher Scientific). The column was maintained at 50°C. Buffer A and Buffer B were 0.1% formic acid in water and 0.1% formic acid in 80% acetonitrile. Peptides were separated on a segmented gradient from 6% to 31% buffer B for 120 min and from 31% to 50% buffer B for 10 min at 200 nl / min. Eluting peptides were analyzed on a QExactive HF mass spectrometer (Thermo Fisher Scientific). Peptide precursor m/z measurements were carried out at 60000 resolution in the 300 to 1800 m/z range. The top ten most intense precursors with charge states from 2 to 7 were selected for HCD fragmentation using 25% normalized collision energy. The m/z values of the peptide fragments were measured at a resolution of 30000 using a minimum AGC target of 8e3 and 55 ms maximum injection time. Upon fragmentation, precursors were put on a dynamic exclusion list for 45 sec.

Bulk proteomics raw data were analyzed using MaxQuant version 1.6.1.0 (Cox & Mann, 2008) and the integrated Andromeda search engine (Cox et al., 2011). Peptide fragmentation spectra were searched against the reviewed and unreviewed sequences of the *Caenorhabditis elegans* reference proteome (Proteome ID UP000001940, downloaded December 2018 from UniProt). Variable modifications included methionine oxidation and protein N-terminal acetylation; cysteine carbamidomethylation was set as a fixed modification. Enzymatic digestion was specified as “Trypsin/P” with “specific” cleavage. The minimum number of peptides and razor peptides required for protein identification was 1, and the minimum number of unique peptides was 0. Protein and peptide spectrum match (PSM) false discovery rates (FDRs) were set to 1% (0.01). The “second peptide” option was enabled. PSM identifications were transferred across raw files using the “Match between runs” feature. Label-free quantification (LFQ) was performed with a minimum ratio count of 2. LFQ intensities were filtered to retain proteins with at least three valid values in at least one experimental group and imputed from a normal distribution with a width of 0.3 and a downshift of 1.8.

### Single worm proteomic sample preparation and analysis

Single adult worms were cleared of bacteria by transferring twice to empty plates and allowing them to crawl for 5 min, then placed into pre-chilled 0.2 mL PCR strip tubes containing 4 µL lysis buffer (0.25% DDM, 125 mM TEAB) and snap-frozen in liquid nitrogen within 3 min. Samples were stored at −80°C until processing. For lysis, worms underwent four freeze–thaw cycles (liquid nitrogen; water at room temperature) followed by sonication in a Bioruptor at 4°C. Digestion was initiated by adding 6 µL chilled PBS pH 7.4 containing a trypsin/Lys-C mix, and samples were incubated at 37°C for 2 hours in a thermocycler. Digests were loaded onto EvoTips for LC–MS/MS.

The Evosep One liquid chromatography system (Bache et al., 2018) was used for sample analysis using the predefined 30 samples per day (30SPD) method. The analytical column was a ReproSil-Pur C18 column (15 cm × 150 µm, 1.9 µm beads; EV1106 Endurance Column, Evosep), operated at 50 °C. Mobile phase A consisted of 0.1% formic acid in water, and mobile phase B was 0.1% formic acid in 100% acetonitrile (ACN).

Peptides were analyzed on a trapped ion mobility spectrometry (TIMS) quadrupole time-of-flight (TOF) mass spectrometer (timsTOF Pro2, Bruker) in data-independent acquisition mode with parallel accumulation, serial fragmentation (diaPASEF). The mass spectrometry scan range was set to m/z 425–1000. Ion accumulation and ramp times in the TIMS module were set to 100 ms. The ion mobility range was defined as 1/K₀ = 0.8–1.25 V·s/cm². The isolation window in the m/z versus ion mobility space was 26 Th. Collision energy was linearly ramped from 20 to 59 eV as a function of ion mobility, ranging from 0.6 to 1.4 V·s/cm².

Raw data were analyzed using Spectronaut (Biognosys AG) against a one-protein-per-gene *C. elegans* reference proteome (UP000001940, August 2022). Methionine oxidation and protein N-terminal acetylation were set as variable modifications, cysteine carbamidomethylation as a fixed modification, and digestion was specified as Trypsin/P with up to two missed cleavages. Protein groups were retained if quantified in all but one sample per comparison group. Missing values were imputed from a down-shifted normal distribution (width 0.3, shift 1.8). Differential protein expression was assessed using limma in R, incorporating sample weights to account for differences in sample quality (Ritchie et al., 2006, 2015).

### Protein stoichiometry analysis

Protein intensities were log_2_-transformed. Samples were trimmed-mean normalized by subtracting, for each sample, the mean logabundance of “always expressed” proteins (proteins quantified in all samples) after excluding proteins in the bottom and top 5% of protein-wise mean abundance.

Ribosomal protein stoichiometry control was assessed within each genotype × age condition by computing pairwise Pearson correlations (r) across samples between all ribosomal proteins, separately for two independent batches (∼15 samples each), and averaging correlation coefficients across batches. To control for abundance-dependent correlation structure (what is expected for proteins with similar abundance), an abundance-matched background set of non-ribosomal proteins was generated per condition by matching the ribosomal proteins’ abundance distribution using quantile binning and random sampling without replacement; background correlations were computed analogously. Correlations were visualized as heatmaps using hierarchical clustering of rows and columns (pheatmap, default settings), and correlation coefficients were summarized by density plots; dashed lines indicate mean r. Differences in correlation distributions relative to the *glp-1* reference within each age group were tested using two-sided Welch’s t-tests on the sets of pairwise correlation coefficients.

### mRNA-Protein Ageing Trajectory Analysis

To identify age-related molecular signatures, we performed Weighted Gene Co-expression Network Analysis (WGCNA (Langfelder & Horvath, 2008)) independently on single-worm transcriptome and proteome datasets. Transcriptomic data were retrieved from the NCBI Sequence Read Archive (BioProject: PRJNA1015633 (Eder et al., 2024b)) while proteomic profiles were generated as described in the preceding sections.

Co-expression networks were constructed using the pyWGCNA package (version 2.2.1), a Python-based implementation of the original WGCNA algorithm (Rezaie et al., 2023). The RNA-sequencing data were preprocessed by selecting 5,000 highly variable genes from the N2 worm aging dataset before module identification. Consensus networks were generated for each dataset across all sampled time points to ensure robust module identification. Modules were characterized based on their correlation with chronological age (days post-maturation). We categorized aging-associated modules using the following criteria based on the correlation coefficient (r) and statistical significance (P < 0.001): Up-regulated: r > 0.6, Down-regulated: r < −0.6, Weakly Up-regulated: 0.2 < r < 0.6, Weakly Down-regulated: −0.6 < r < −0.2, No Change (nc): All remaining modules that did not meet the significance or correlation thresholds.

To compare expression dynamics across omics layers, individual gene and protein expression levels were normalized using Z-score transformation across the time axis. This allowed for the direct comparison of relative abundance changes during the aging process. The overlap of gene membership between transcriptomic and proteomic modules was visualized using Sankey diagrams, facilitating the identification of coordinated or anti-correlated molecular signatures between the two layers

### Senescence-associated β galactosidase staining (SA-≥-gal)

SA-β-galactosidase activity was assessed in synchronized adult worms as a marker of age-associated lysosomal β-galactosidase activity. Equal numbers of synchronized worms per condition (∼100) were collected and mechanically permeabilized by freeze–thaw before fixation to facilitate substrate access (Duerr, 2013). Worms were then subjected to chromogenic β-galactosidase staining with X-Gal under standard conditions, following methods described in (Nonninger et al., 2025; Venz et al., 2024), which document in vivo β-galactosidase activity in *C. elegans*.

After staining, samples were imaged on a Zeiss Axio Imager Z2 microscope using consistent acquisition settings. All biological replicates for a given experiment were imaged on the same day to minimize technical variability. The stained signal was quantified from bright-field images using Fiji.

### Fluorescence microscopy and image analysis

Bright-field, differential interference contrast (DIC), and fluorescence images were acquired using either a Zeiss Axio Imager Z1 microscope or a Leica M205 FA stereomicroscope, depending on the experimental application. High-resolution DIC and fluorescence imaging were performed on a Zeiss Axio Imager Z1 equipped with a Colibri 7 LED light source, Axio Cam Mono 506 and AxioCam ICc5 cameras, and standard filter sets for GFP (Set 38 HE) and TR (Set 45). Image acquisition was controlled using Zeiss ZEN software (v2.3.69.1017). Lower-magnification fluorescence imaging was performed using a Leica M205 FA stereomicroscope equipped with a DFC3000G monochrome camera and an EL6000 external light source, operated with Leica Application Suite X software (v3.7.5.24914).

Nucleolar size measurements were performed on *fib-1::neongreen* fluorescence images acquired on the Zeiss Axio Imager Z1 using a 100× objective. Synchronized worms were immobilized with 0.01% sodium azide and imaged after cessation of movement. Nucleolar area was manually quantified in the head region using the freehand selection tool in Fiji. For *mrps-16::mKate* experiments, fluorescence images were acquired using the Leica stereomicroscope and analyzed in Fiji.

Confocal imaging was performed on a Leica TCS SP8 confocal microscope equipped with a white light laser and a ×63/1.4 NA oil immersion objective, with HyD detectors for fluorescence and a PMT detector for bright-field imaging. Image acquisition was performed using LAS X Life Science software. For *mKate::popl-1* experiments, nuclear fluorescence intensity was quantified from confocal images using Fiji based on manually segmented nuclear regions in the head region.

### SUnSET assay and Immunoblotting

Global protein synthesis was assessed by puromycin incorporation (SUnSET assay). Synchronized worms were collected, washed extensively in S basal buffer (Stiernagle, 2006), and incubated in liquid culture containing heat-killed bacteria corresponding to each RNAi condition. Approximately 2,000 worms per condition were incubated in S medium at 20°C with shaking (150 rpm), and puromycin was added to a final concentration of 1 mM for 4 h. Worms were then washed to remove excess puromycin, pelleted, flash-frozen, and stored at −80°C.

Frozen pellets were lysed in CelLytic lysis buffer supplemented with protease inhibitors by repeated freeze–thaw cycles and sonication. Lysates were clarified by centrifugation, and protein concentration was determined by BCA assay. Equal amounts of protein were resolved by SDS–PAGE on stain-free Criterion TGX gels and transferred to nitrocellulose membranes. Membranes were probed with an anti-puromycin monoclonal antibody (clone 3RH11) followed by HRP-conjugated secondary antibody, and signals were detected by chemiluminescence using a ChemiDoc MP imaging system.

### Polysome Profiling

Polysome profiling was performed essentially as described previously (Derisbourg et al., 2021), with minor adaptations. All steps were carried out at 4°C or on ice in the presence of cycloheximide to preserve ribosome–mRNA complexes.

Synchronized adult worms (5,000–10,000 per condition) were harvested from plates, washed extensively in M9 buffer, and pelleted. Pellets were either processed immediately or flash-frozen in liquid nitrogen and stored at −80°C. Worms were lysed in polysome lysis buffer containing Igepal, sodium deoxycholate, DTT, RNAse inhibitor, protease inhibitors, and cycloheximide using a Dounce homogenizer. Lysates were clarified by centrifugation, and protein concentration was determined by BCA assay. Equal amounts of lysate (typically 500–1,000 µg total protein) were loaded onto linear 10-60% (w/v) sucrose gradients. Samples were resolved by ultracentrifugation in an SW40Ti rotor at 39,000 rpm for 3 h at 4°C. Gradients were analyzed by continuous UV absorbance monitoring at 254 nm using a Biocomp gradient fractionation system.

For quantification, absorbance traces were plotted using GraphPad Prism. The area under the curve corresponding to monosome and polysome fractions was quantified using Fiji to assess relative ribosome distribution across conditions

### Ribosome Sequencing: sample preparation, library preparation, and sequencing

Ribosome profiling was performed with minor adaptations from established *C. elegans* protocols. Briefly, ∼10,000 synchronized adult worms were collected, washed in M9 buffer, flash-frozen, and thawed in the presence of cycloheximide (1 mM). Worms were lysed mechanically in polysome lysis buffer (Tris-HCl 20 mM, KCl 140 mM, MgCl₂ 1.5 mM, NP-40 0.5 %, sodium deoxycholate 1%, DTT 1 mM, PMSF 1 mM, and cycloheximide 1 mM). Clarified lysates were treated with RNAse I for 1 h at room temperature to generate ribosome-protected fragments.

Digested lysates were resolved on linear 15–50% sucrose gradients by ultracentrifugation (SW40Ti rotor, 39,000 rpm, 2 h, 4°C) and flash-frozen. RNA was extracted by proteinase K digestion followed by acid phenol–chloroform extraction and ethanol precipitation. Purified RNA was further cleaned using the Zymo RNA Clean & Concentrator-5 kit and submitted as dry pellets for library preparation. Total RNA samples used for normalization were processed in parallel using the RNA-seq protocol described above.

RNA samples were treated with T4 Polynucleotide kinase for 90 min at 37°C. Afterwards, samples were run on denaturing polyacrylamide gels (15% polyacrylamide TBE-urea gels, Thermo Fisher) and bands corresponding to 25 to 35nt were excised. RNA was extracted from the gel, purified, and precipitated. The RNA pellet was resuspended in nuclease-free water, and RNA concentrations were measured with Qubit (Qubit RNA HS, Thermo Fisher). Libraries were prepared with the Lexogen small RNA Library Prep kit (16 PCR cycles). Next, library validation and quantification (Agilent Tape Station) was performed, followed by pooling of equimolar amounts of library. The pools were then quantified using the Peqlab KAPA Library Quantification Kit and the Applied Biosystems 7900HT Sequence Detection System and sequenced on an Illumina NovaSeq6000 sequencing instrument with an SR100 protocol aiming for 100 million clusters per sample.

Riboseq reads were trimmed with Flexbar (v3.5) and mapped to the *Caenorhabditis elegans* reference genome (Ensembl 110) using bowtie (v1.2.3). After removal of reads mapping to rRNA or tRNA, reads were mapped to the same reference genome and transcriptome using START (v2.7.3a). P-site offset, 3-nt periodicity, codon usage, and meta-profiles were calculated with riboWaltz (v2.0). Pausing sites were identified with Paus,ePred, and a 5’ tag profile was generated for genes with a pausing site using Rfeet. Actively translated ORFs were found using Pythonthon library RiboCode (v1.2.15). Differentially translated genes were identified using DESeq2 (v1.24.0). Translation efficiency was calculated using riborex (v2.4.0).

### SDS-Insoluble aggregation analysis

All *glp-1* mutant strains were maintained at 15°C for at least two generations after thawing. For synchronization, ∼200 gravid adults were placed on NGM plates seeded with concentrated OP50 (20×) and incubated at 15°C for 72 h. Gravid adults were collected in S-basal, washed three times to remove bacteria, and embryos were isolated by alkaline hypochlorite treatment. Eggs were washed ≥5 times in M9 to remove bleach and hatched for 24 h at 25°C in M9 with rotation to obtain synchronized L1 larvae. L1s were plated on NGM seeded with concentrated HT115 (5×) expressing either scrambled control RNAi or RNAi targeting the gene of interest and maintained at 25°C. Animals were transferred to 20°C at day 1 and to fresh RNAi plates at day 2 of adulthood and harvested at day 5 by washing plates with S-basal, followed by ≥3 washes by gravity settling and aspiration. Pellets were flash-frozen in a dry ice/ethanol bath.

Frozen worm pellets were thawed in detergent-free lysis buffer (20 mM Tris base, pH 7.4, 100 mM NaCl, 1 mM MgCl_2_)containing protease inhibitors and lysed by water-bath sonication at 4°C. Lysis efficiency was verified by light microscopy. Lysates were clarified by centrifugation, and protein concentration was determined by BCA. Insoluble proteins were enriched by pelleting the lysate corresponding to 1 mg total protein (20,000 g, 15 minutes at 4°C). The pellet was extracted three times in 1% SDS lysis buffer (20,000 g, 15 minutes; room temperature). The resulting SDS-insoluble pellet was re-solubilized in 70% formic acid with sonication, dried in a vacuum concentrator to remove formic acid. The dried pellet was resuspended in 1× LDS sample buffer and heated at 95°C for 10 minutes. Samples were resolved by SDS–PAGE (4–12% NuPAGE Bis-Tris), stained with SYPRO Ruby (Invitrogen), imaged on a ChemiDoc system (Bio-Rad), and quantified in ImageJ (NIH).

## Acknowledgements

We thank the CGC (University of Minnesota) and the Japanese National Biosource Project for providing *C. elegans* strains and the Cologne Centre for Genomics for RNA-sequencing and Ribosome sequencing. F. Metge, A. Iqbal, M. Duhamel, and J. Boucas from the MPI-AGE Bioinformatics, C. Kukat and M. Kirchner from MPI-AGE Imaging, and X. Li, I. Atanassov, and T. Colby from MPI-AGE Proteomics Cores for technical support. We also thank S. Kreuz for valuable comments on the manuscript. T. Popkes von Oepen was supported by the International Max Planck Research School, Cologne Graduate School of Ageing Research, and the project was funded through the Max Planck Gesellschaft. The project leading to this application has received funding from the European Research Council (ERC) under the European Union’s Horizon 2020 research and innovation programme (grant agreement No 834259). This work was also supported by the Max Planck Society, Germany.

## Author contributions

V.E.M-M. performed experiments, analyzed data, helped write the manuscript, prepared figures, and provided intellectual input. T.P.v O. helped conceive and design the study, performed experiments, analyzed data, and helped prepare figures. T.S.S performed experiments and analyzed data. Y.P. performed the RNA and protein coupling analysis. F.L. performed the ribosomal stoichiometry analysis. W.H. provided the N2 ageing single worm proteomics data. L.S.J. gave technical support. E.A. performed SDS-insoluble in-gel quantifications. G.J.L. helped design experiments and provided intellectual input. A.B. contributed to the analysis and provided intellectual input. A.A. conceived the project, secured funding, and wrote the manuscript. All authors discussed the results and commented on the manuscript.

## Supplemental information

**Figure S1.**
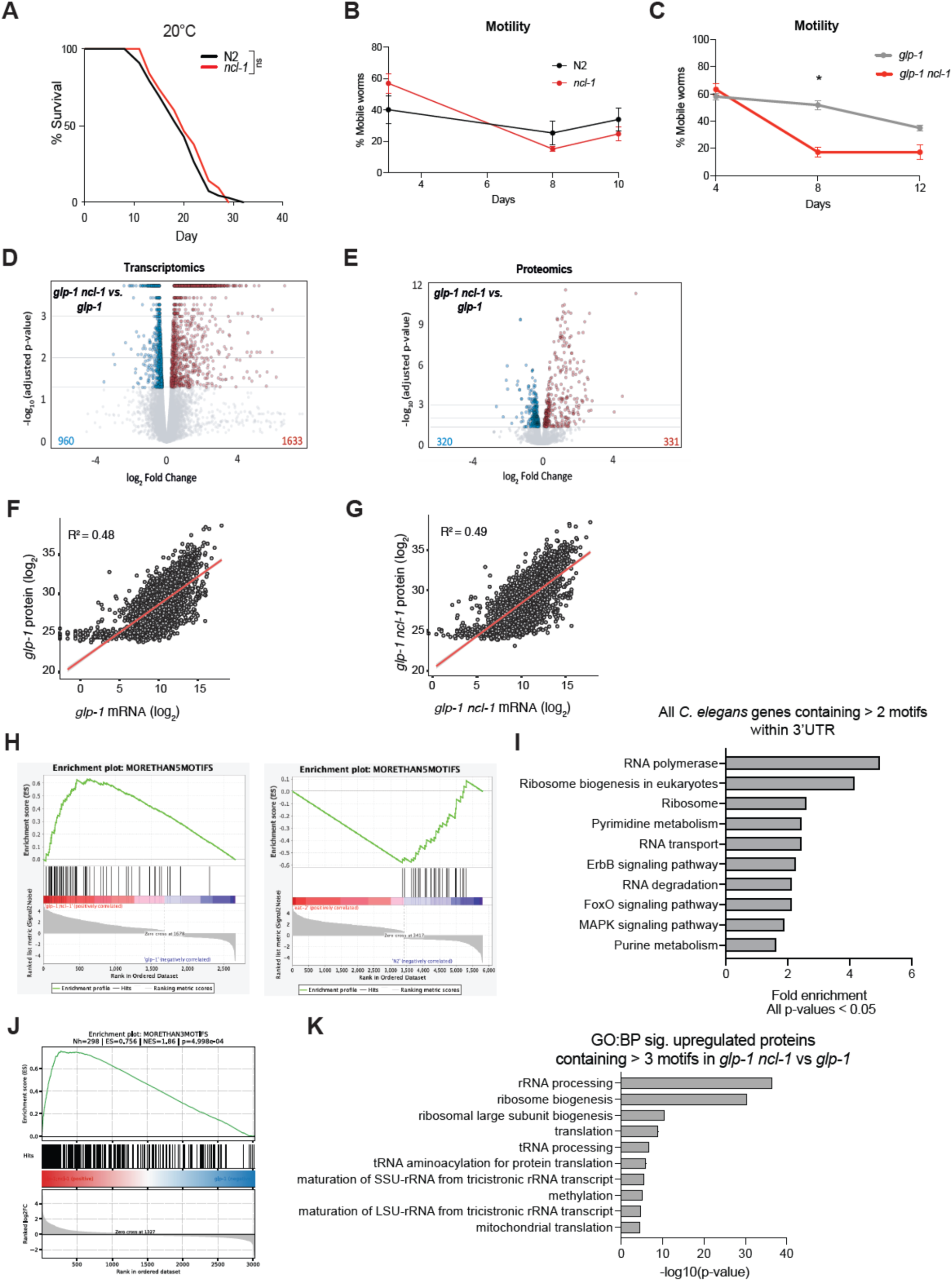
Transcriptomic, proteomic, and motif-enrichment analyses associated with *ncl-1* loss. **A**. Lifespan analysis of N2 and *ncl-1 (e1942)* worms reared at 20°C since egg lay. (n>100, log-rank Mantel-Cox test) Data on Table 1. **B**. Age-dependent mobility assay of *glp-1* and *glp-1 ncl-1* worms. Mobility was quantified as the percentage of worms leaving a **⌀** 1 cm circle after 2 minutes. Mean of 5 technical replicates with 20 worms/replicate. (2-way ANOVA post hoc Tukey’s multiple comparisons test, mean ± SEM). **C.** Percentage of mobile *glp-1 (e2141)* and *glp-1 (e2141) ncl-1 (e1942)* worms leaving a **⌀** 1 cm circle after 2 minutes. Mean of 5 technical replicates with 20 worms/replicate. *glp-1 ncl-1* vs *glp-1* at day 8 are significantly less mobile (p=0.0253). (2-way ANOVA post hoc Tukey’s multiple comparisons test, mean ± SEM). **D**. Volcano plot showing differential gene regulation in *glp-1 ncl-1* mutants compared to *glp-1* at day 1. Logfold change values are plotted against the adjusted-logp-values. Each dot represents a gene, while upregulated candidates with a p. adj <0.05 are labeled red and downregulated are labeled blue. The total number of significantly up- or downregulated candidates is depicted in the bottom corners of each graph. **E**. Volcano plots showing differential protein regulation in *glp-1 ncl-1* mutants compared to *glp-1* at day 1. Logfold change values are plotted against the adjusted-logp-values. Each dot represents a protein, while upregulated candidates with a p. adj <0.05 are labeled red and downregulated are labeled blue. The total number of significantly up- or downregulated candidates is depicted in the bottom corners of each graph. F. Correlation plot of mRNA and protein abundance of log2-transformed raw counts of mRNA and proteins plotted against each other in *glp-1* worms at day 1. G. Correlation plot of mRNA and protein abundance of log2-transformed normalized counts of mRNA and proteins plotted against each other in *glp-1 ncl-1* worms at day 1 **H**. Genes ranked by their correlation with the *glp-1 ncl-1* expression profile compared to *glp-1* (left) and another longevity mutant, *eat-2(ad465)*, compared to N2 (right). The enrichment score (ES) (green) shows that TTGTT-rich transcripts (black ticks) containing more than 5 motifs are concentrated toward the positively correlated end of the ranked list in *gpl-1 ncl-1*, indicating enrichment among transcripts upregulated in the *ncl-1* loss-of-function mutant compared to the *glp-1* controls. The lower panel shows the ranking metric across the ordered gene list, with the “zero cross” marking the point where the cumulative enrichment score transitions. *eat-2* mutants do not show an enrichment of UUGUU motifs in the top-ranked terms compared to N2. I. Kegg pathway enrichment analysis of all genes in the *C. elegans* genome containing more than two TTGTT motifs downstream of the stop codon. **J**. Proteins ranked by log₂ fold change in *glp-1 ncl-1* versus *glp-1* proteomic profiles. The enrichment score (ES, green) shows that proteins containing more than three post-stop UUGUU motifs (black ticks) are significantly enriched toward the upregulated end of the ranked proteome in *glp-1 ncl-1* mutants. The lower panel shows the ranked log₂ fold change metric across the ordered protein list, with the zero-cross indicating the transition point between up- and downregulated proteins. Enrichment significance was assessed by permutation testing (NES and p-value shown). K. GO Biological Process enrichment analysis of significantly upregulated proteins in glp-1 ncl-1 versus glp-1 that contain more than three post-stop UUGUU motifs. Enriched terms highlight ribosome biogenesis, rRNA processing, and translation-related processes (DAVID analysis; p < 0.05).

**Figure S2.**
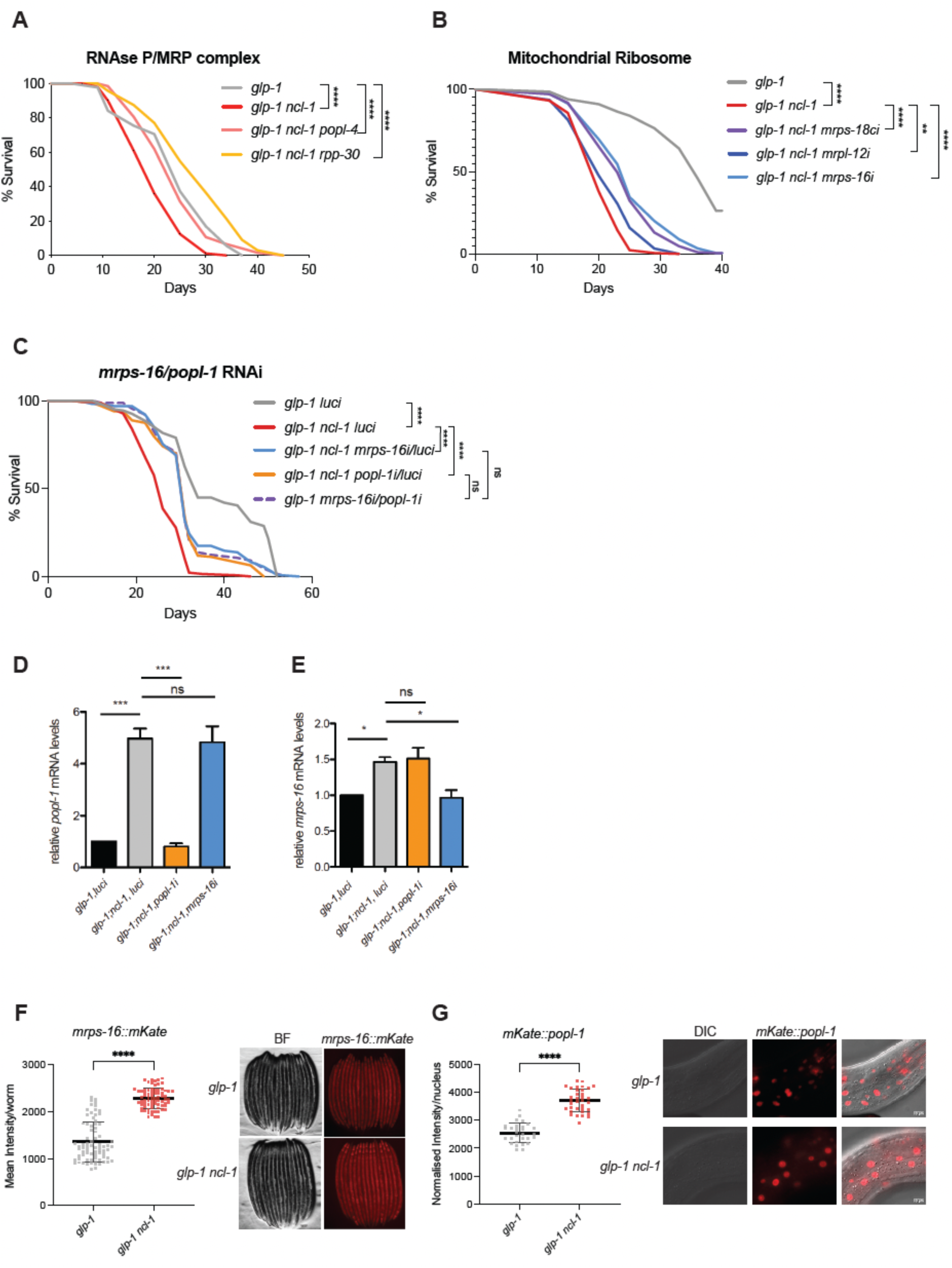
RNAi targeting mitochondrial ribosome and RNAse P/MRP components restores longevity to *glp-1 ncl-1* double mutants. **A**. Lifespan analysis of *glp-1* and *glp-1 ncl-1* worms on *luci* control, or RNAse P/MRP components, *popl-4i* and *rpp-30I*, which show lifespan extension. All curves are statistically significant (p <0.0001, n>100, log-rank Mantel-Cox test). Data from all replicates are shown in Table 1. **B**. Lifespan analysis of *glp-1* and *glp-1 ncl-1* worms on *luci* control, or on RNAi of mitochondrial ribosome subunits, *mrps-5ci*, and *mrpl-45i*, showing lifespan extension. All curves are statistically significant (p <0.0001), except *glp-1 ncl-1 mrps-5i* compared to *glp-1 ncl-1* (p =0.0005, n>100, log-rank Mantel-Cox test). Data from all replicates are shown in Table 1. **C**. Lifespan analysis of *glp-1* on *luci* control and *glp-1 ncl-1* worms on *luci*, *mrps-16i*/*luci* double RNAi, *popl-1*i/*luci* double RNAi, and *mrps-16i/ popl-1i* double RNAi, showing extension of lifespan. All curves are statistically significant (p <0.0001), except *glp-1 ncl-1 mrps-16i/popl-1i* compared to *glp-1 ncl-1 mrps-16i/luci* and *glp-1 ncl-1 mrps-16i/popl-1i* compared to *glp-1 ncl-1 popl-1i/luci*, which are not significant (n>100, log-rank Mantel-Cox test), showing no additive effect of *popl-1i* / *mrps-16i* double RNAi. Data from all replicates are shown in Table 1. D. qRT-PCR analysis of *popl-1* mRNA in *glp-1* and *glp-1 ncl-1* mutants on luciferase, *mrps-16i* or *popl-1i* at day 1 of adulthood. The level of *popl-1* mRNA is significantly increased in *glp-1 ncl-1* on luciferase compared to *glp-1*, and down in *glp-1 ncl-1* on *popl-1i* compared to *glp-1 ncl-1* on *luci* (p <0.001). (n=3, One-way ANOVA post hoc Dunnett’s multiple comparison test, mean ± SEM). E. qRT-PCR against *mrps-16* mRNA in *glp-1* and *glp-1 ncl-1* mutants on luciferase, *mrps-16i* or *popl-1i* worms at day 1 of adulthood. The level of *popl-1* mRNA is significantly increased in *glp-1 ncl-1* on luciferase compared to *glp-1*, and down in *glp-1 ncl-1* on *mrps-16i* compared to *glp-1 ncl-1* on *luciferase* (p <0.05, n=3, One-way ANOVA post hoc Dunnett’s multiple comparison test, mean ± SEM). F. Mean fluorescence intensity of *mrps-16* endogenously tagged with mKate by CRISPR-Cas9 at the L4 larval stage in *glp-1* and *glp-1 ncl-1* worms. The level of *mrps-16::mKate* is significantly increased in *glp-1 ncl-1* on luciferase compared to *glp-1* (p <0.0001). Each dot represents a worm. (n=3, One-way ANOVA post hoc Dunnett’s multiple comparison test, mean ± SD) G. Mean fluorescence intensity of *popl-1* endogenously tagged with mKate by CRISPR-Cas9 at day 1 of adulthood in nuclei from *glp-1* and *glp-1 ncl-1* worms on luciferase. The level of *mKate::popl-1* is significantly increased in *glp-1 ncl-1* on luciferase compared to *glp-1*, and significantly down in *glp-1 ncl-1* on *popl-1i* compared to *glp-1 ncl-1* on *luciferase* (p <0.0001). Each dot represents a nucleus. (n=3, One-way ANOVA post hoc Dunnett’s multiple comparison test, mean ± SD)

**Figure S3.**
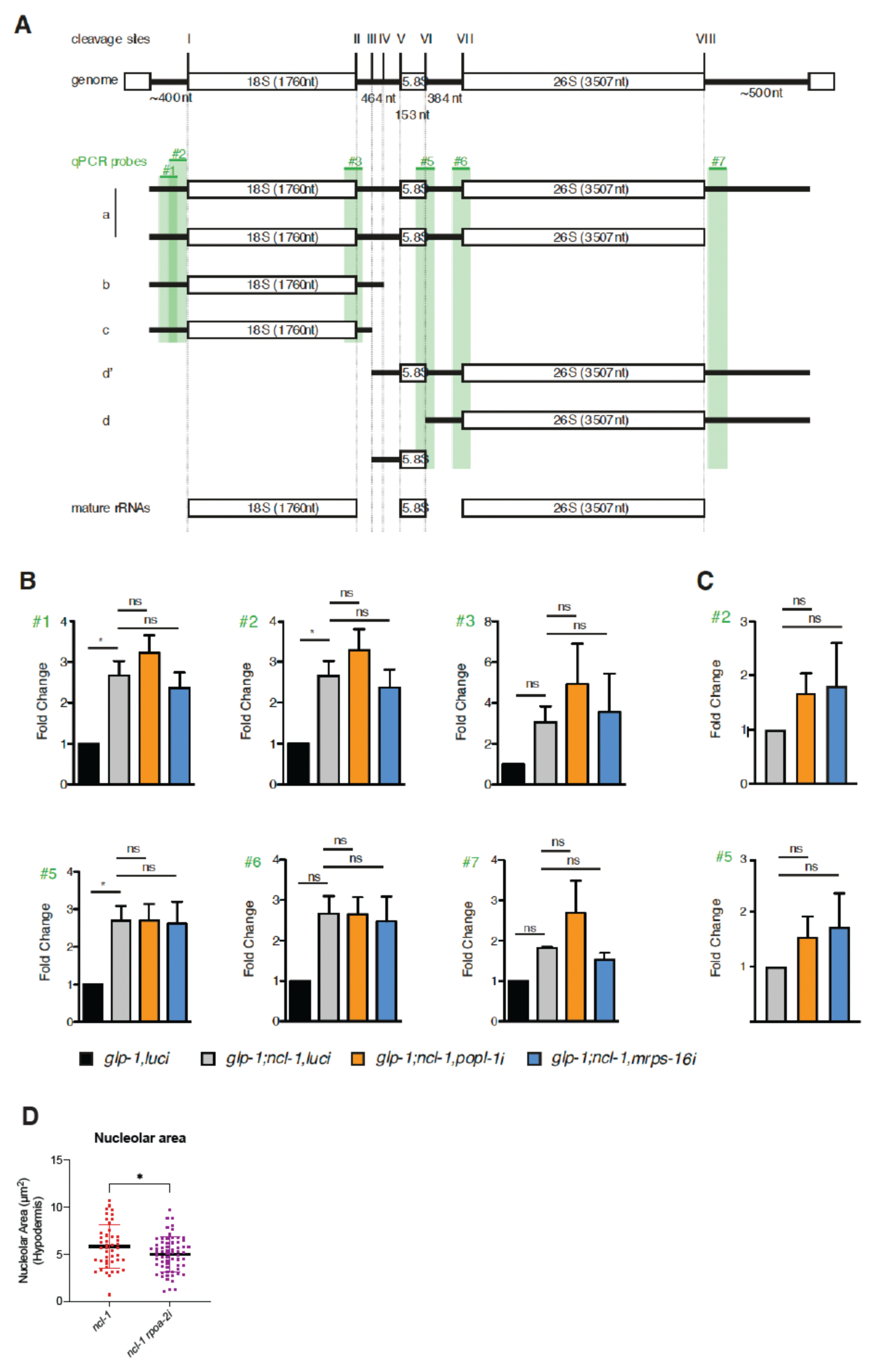
*ncl-1* loss drives accumulation of precursor rRNA and nucleolar enlargement through increased rRNA precursor transcription **A**. Schemaäc representaäon of rRNA processing steps and intermediates, indicaäng the posiäons of qRT–PCR probes (green) targeäng precursor rRNA regions. Probes do not detect mature rRNA. Adapted from Saijou et al. (2004). **B**. qRT-PCR against the probes depicted in A. showing increased precursors in *glp-1 ncl-1* on luciferase control, *mrps-16i*, and *popl-1i* compared to *glp-1* on luciferase at day 1 of adulthood. Measurements normalized to *snb-1* mRNA. (n=3, One-Way ANOVA post hoc Dunnett’s multiple comparison test, mean ± SEM). **C**. qRT-PCR against the probes depicted in A, showing increased precursors in *glp-1 ncl-1* on luciferase control, *mrps-16i*, and *popl-1i* compared to *glp-1* on luciferase at day 6 of adulthood. Measurements normalized to *snb-1* mRNA. (n=3, One-Way ANOVA post hoc Dunnett’s multiple comparison test, mean ± SEM). D. Quantification of nucleolar area (m^2^) measured in hypodermal cells of *ncl-1 (e1942)* worms fed luciferase control (red) or *rpoa-2i* (purple) at day 1 of adulthood. *RPOA-2i* significantly reduces nucleolar size (p=0.0335). Each dot represents a nucleolus. (n=40-50 nucleoli from 12-15 worms, Unpaired t-test, mean ± SD).

**Figure S4.**
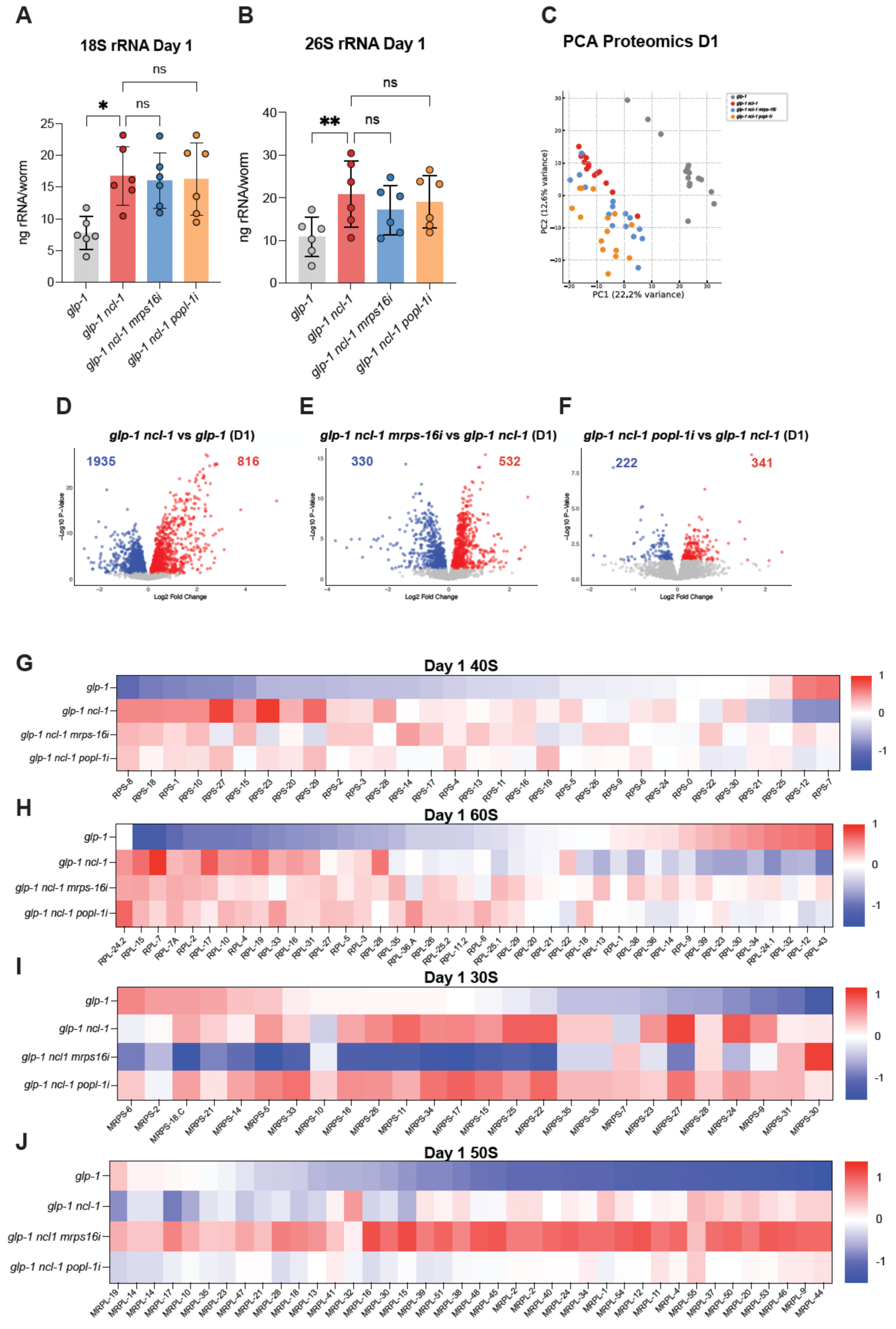
*ncl-1* life-extending suppressors maintain rRNA abundance and ribosomal protein levels over age. A. Mature 18S rRNA levels were measured at day 1 of adulthood. *glp-1 ncl-1* compared to *glp-1* have increased 18S rRNA (p =0.011). Treatment with *mrps-16i* or *popl-1i* does not significantly alter 18S rRNA levels relative to *glp-1 ncl-1*. Each dot represents a biological replicate of 100 worms. (n=6, Friedman ANOVA post hoc Dunn’s multiple comparisons test, mean ± SEM). B. Mature 26S rRNA levels were measured at day 1 of adulthood. *glp-1 ncl-1* compared to *glp-1* have increased 26S rRNA (p =0.0024), with no significant change upon *mrps-16i* or *popl-1i* treatments relative to *glp-1 ncl-1*. Each dot represents a biological replicate of 100 worms. (n=6, Friedman ANOVA post hoc Dunn’s multiple comparisons test, mean ± SEM). C. PCA of single-worm LC-MS/MS proteomic data from *glp-1* (grey), *glp-1 ncl-1* (red), *glp-1 ncl-1* on *mrps-16i* (blue), and *glp-1 ncl-1* on *popl-1i* (orange) at day 1 of adulthood. Each point corresponds to one worm (n=16). The clustering indicates that *glp-1* and *glp-1 ncl-1* separate primarily along PC1, while *glp-1 ncl-1* and *glp-1 ncl-1* on *mrps-16i* or *popl-1i* distribute on PC2. **D**. Volcano plots of differentially expressed proteins in day 1 *glp-1 ncl-1* worms compared to *glp-1*, *glp-1 ncl-1* on *mrps-16i*, compared to *glp-1 ncl-1* **E**., and *glp-1 ncl-1* on *popl-1i* compared to *glp-1 ncl-1* **F**. Blue dots indicate significantly downregulated and red significantly upregulated proteins (adjusted p < 0.05). Logfold changes in normalized expression counts. G. Average Z-scores of all the cytosolic ribosomal proteins belonging to the 40S (G) and 60S (H) subunits detected in the single worm proteomics at day 1 in *glp-1*, *glp-1 ncl-1*, *glp-1 ncl-1 mrps-16i*, or *popl-1i*. I. Average Z-scores of all the mitochondrial ribosomal proteins from the 30S (I) and 50S (J) subunits detected in the single worm proteomics at day 1 in *glp-1*, *glp-1 ncl-1*, *glp-1 ncl-1 mrps-16*, or *popl-1i*. For each protein, the mean across all samples was subtracted and divided by the standard deviation. 0 indicates expression equal to the protein’s average, positive values (red), higher-than-average, and negative (blue), lower-than-average.

**Figure S5.**
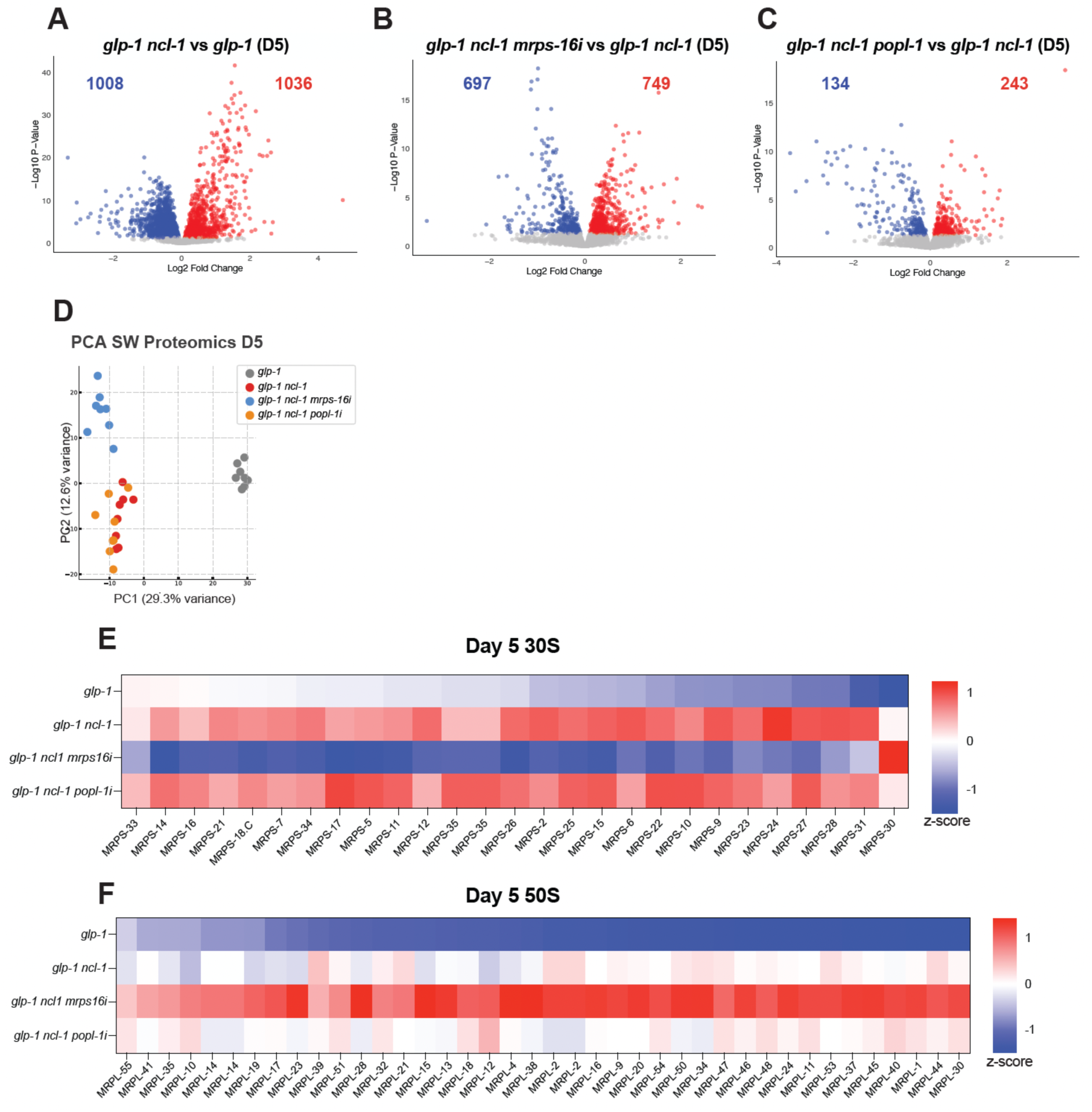
Lifespan-extending RNAi treatments remodel the day 5 proteome **A**. Volcano plots of differentially expressed proteins from single-worm proteomics in day 5 *glp-1 ncl-1* worms compared to *glp-1*, *glp-1 ncl-1* on *mrps-16* RNAi, compared to *glp-1 ncl-1* **B**., and *glp-1 ncl-1* on *popl-1i* compared to *glp-1 ncl-1* **C**. Blue dots indicate significant downregulation and red significant upregulation (p adj. < 0.05). Logfold changes in normalized expression counts. D. PCA of single-worm proteomic data from *glp-1* (grey), *glp-1 ncl-1* (red), *glp-1 ncl-1* on *mrps-16i* (blue), and *glp-1 ncl-1* on *popl-1i* (orange) at day 1. Each point corresponds to one worm (n=8). The analysis was carried out on normalized protein expression counts, first and second principal components (PC1, PC2) are shown. The clustering indicates that *glp-1* and *glp-1 ncl-1* separate primarily along PC,1 and *glp-1 ncl-1* and *glp-1 ncl-1* on *mrps-16i* distribute along PC2. E. Average Z-scores of all the mitochondrial RPs from the 30S (E) and 50S (F) subunits detected in single worm proteomics at day 5 in *glp-1*, *glp-1 ncl-1*, *glp-1 ncl-1 mrps-16i*, or *popl-1i*. For each protein, the mean across all samples was subtracted and divided by the standard deviation. 0 indicates expression equal to the protein’s average, positive values (red), higher-than-average, and negative (blue), lower-than-average.

**Figure S6.**
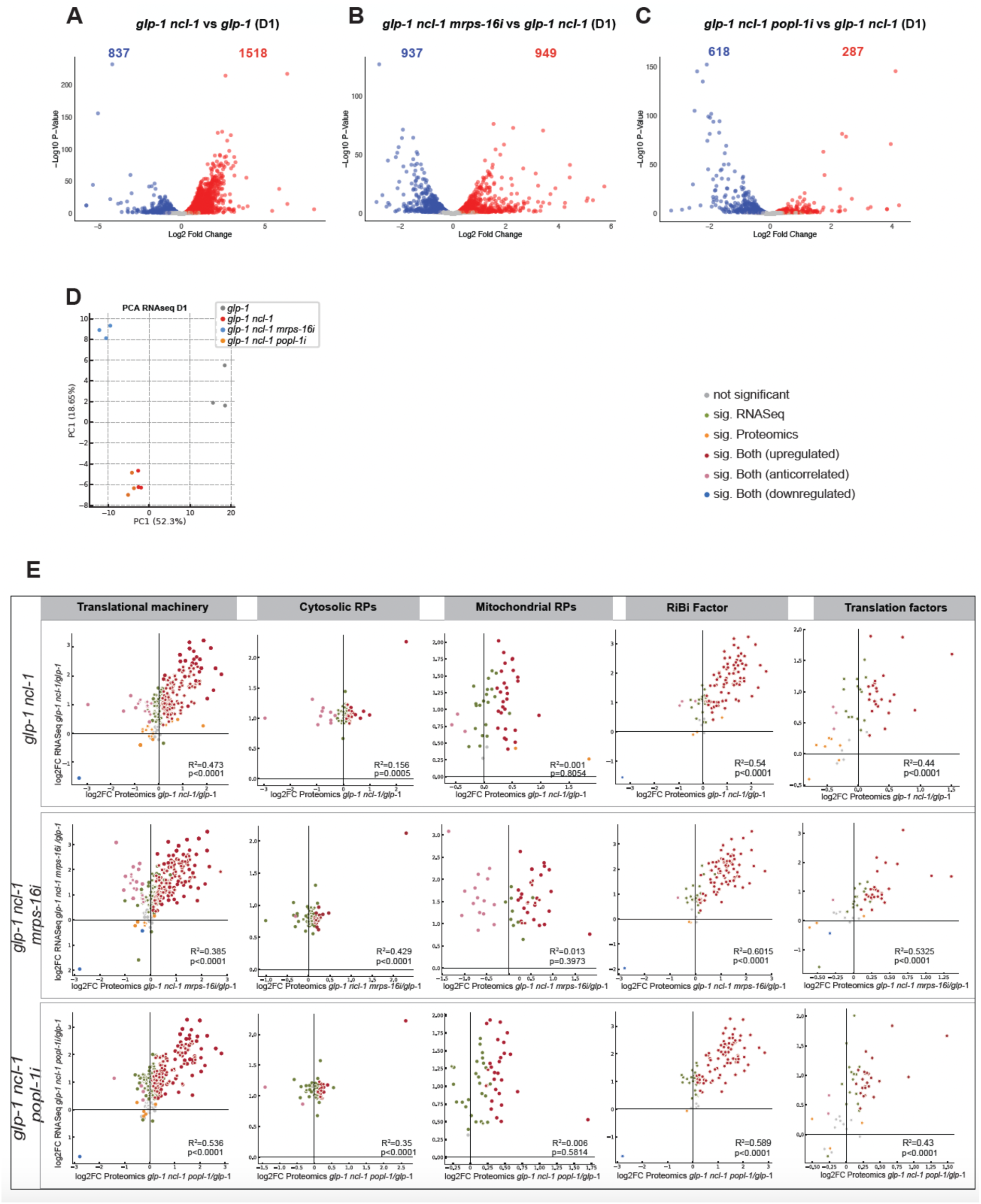
RNAi suppression of *ncl-1* restores transcript-protein coordination within the ribosome biogenesis and translational machinery. A. Volcano plots of differentially expressed transcripts in day 1 *glp-1 ncl-1* worms compared to *glp-1*, *glp-1 ncl-1* on *mrps-16i* compared to *glp-1 ncl-1* B., and *glp-1 ncl-1* on *popl-1i* compared to *glp-1 ncl-1* C. Blue dots indicate significant downregulation and red significant upregulation (adjusted p < 0.05). D. PCA of RNA-seq data from *glp-1* (grey), *glp-1 ncl-1* (red), *glp-1 ncl-1* on *mrps-16i* (blue), and *glp-1 ncl-1* on *popl-1i* (orange) at day 1. Each point corresponds to a biological replicate (n=3). The analysis was carried out on normalized gene-expression counts, first and second principal components (PC1, PC2) are shown. Clustering indicates that *glp-1* and *glp-1 ncl-1* separate primarily along PC1, while *glp-1 ncl-1* and *glp-1 ncl-1* on *mrps-16i* distribute along PC2. **J**. Co-regulation of all the detected transcripts and proteins, logfold changes of the RiBi factors and translation factors (as categorized by GO: BP) in day 1 worms. Grey represents genes that are not significant in both datasets, green is significant only in RNAseq, orange only in proteomics, red is significantly upregulated in both datasets, pink is significant in both datasets but in opposite direction, and blue is significantly downregulated in both datasets. (Pearson R^2^ values; p values indicated in the figure) Proteins showing anticorrelation: RPS-7, RPS-12, RPL-12, RPL-23, RPL-24.1, RPL-30, RPL-32, RPL-34, RPL-39, and RPL-43

**Figure S7.**
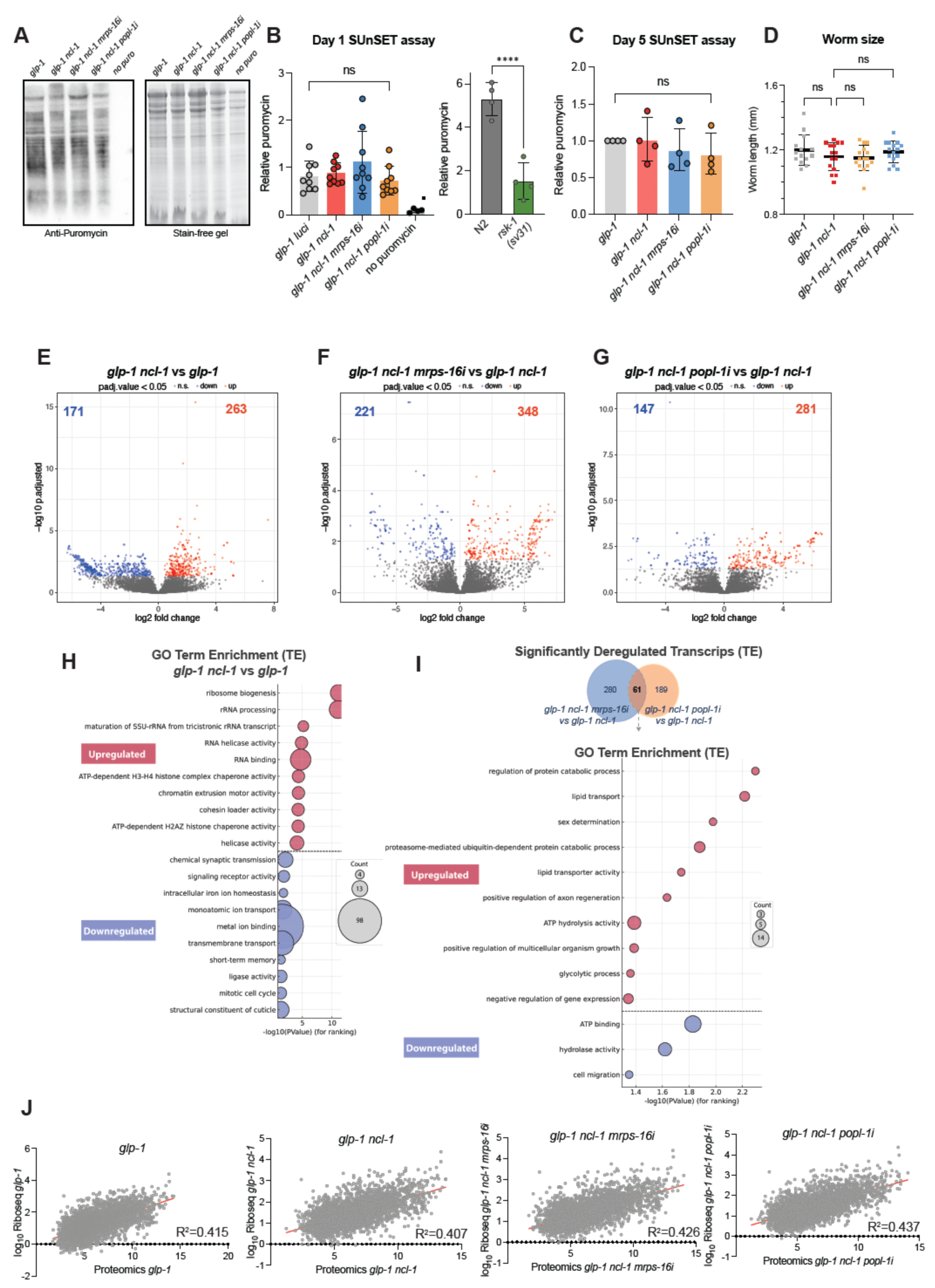
Loss of *ncl-1* uncouples ribosome biogenesis from bulk protein synthesis and redistributes translational output. **A.** Representative SUnSET-based western blot of global protein synthesis in *glp-1 ncl-1*, *glp-1 ncl-1 mrps-16*, or *popl-1i* at day 1 of adulthood. Lysates from worms of the indicated genotypes subjected to the SUnSET assay were separated by SDS-PAGE, transferred to membranes, and probed with anti-puromycin antibody to detect puromycin-labeled nascent polypeptides (left panel). The corresponding stain-free gel (Bio-Rad) image is shown on the right as a loading control for total protein. **B**. Quantification of the SUnSET signal at day 1 of adulthood. Background-subtracted band intensities were normalized to total protein per lane and expressed as fold change relative to the mean of the *glp-1* control group. Each dot represents a biological replicate. (n=10, Friedman one-way ANOVA post hoc Dunnett’s multiple test, mean ± SEM). Control quantification of the SUnSET-based western blot of N2 and *rsks-1 (sv31)* mutants for control of the assay’s sensitivity (p <0.0001). Each dot represents a biological replicate. (n=4, Friedman one-way ANOVA post hoc Dunnett’s multiple test, mean ± SEM). C. Quantification SUnSET signal of *glp-1*, *glp-1 ncl-1*, *glp-1 ncl-1 mrps-16i* and *glp-1 ncl-1 popl-1i* worms at day 5 of adulthood. Each dot represents a biological replicate. (n=4, Friedman one-way ANOVA post hoc Dunnett’s multiple test, mean ± SEM). D. Body length (mm) of day 1 adult *glp-1*, *glp-1 ncl-1*, *glp-1 ncl-1 mrps-16i*, and *glp-1 ncl-1 popl-1i* worms. Each dot represents an individual worm. (n=15, One-way ANOVA post-hoc Turkey’s multiple comparisons test, mean ± SD). **E**. Volcano plots of differentially expressed normalized Ribo-Sequencing transcript counts in day 1 *glp-1 ncl-1* worms compared to *glp-1*, *glp-1 ncl-1* on *mrps-16i* compared to *glp-1 ncl-1* **F**., and *glp-1 ncl-1* on *popl-1i* compared to *glp-1 ncl-1* **G**. Blue dots indicate significant downregulation and red significant upregulation (adjusted p < 0.05). Logfold changes in Ribo-sequencing counts. H. Top 10enriched GO biological process (GO: BP) in the significantly deregulated proteins (p<0.05) in *glp-1 ncl-*compared to *glp-1*. I. Venn Diagram showing 61shared differentially translated transcripts (DETs) in *glp-1 ncl-1* worms treated with *mrps-16i* or *popl-1i* compared to *glp-1 ncl-1*. Top enriched GO: BP and GO molecular function (GO: FM) terms from the overlapped DETs. J. Correlation plots of Ribo-Seq and protein abundance (LC-MS/MS) at day 1 of adulthood in *glp-1*, *glp-1 ncl-1*, *glp-1 ncl-1 mrps-16i*, and *glp-1 ncl-1 popl-1i* worms. Axes showlog10-transformedd raw counts of transcripts and proteins plotted against each other. K. Correlation of logfold changes in the single worm proteomics of *ncl-1* loss of function compared to N2, and the insoluble proteins from *ncl-1* mutants compared to N2. All samples were measured at day 5. Each dot represents a protein. Blue dots represent significantly downregulated proteins in both data sets, red significantly upregulated in both data sets, green significantly regulated in opposite directions, and grey not significant in one of the sets. The grey dashed line denotes x = y, and the solid line the regression line, R^2^ = 0.0642 (p <0.0001).

**Figure S8.**
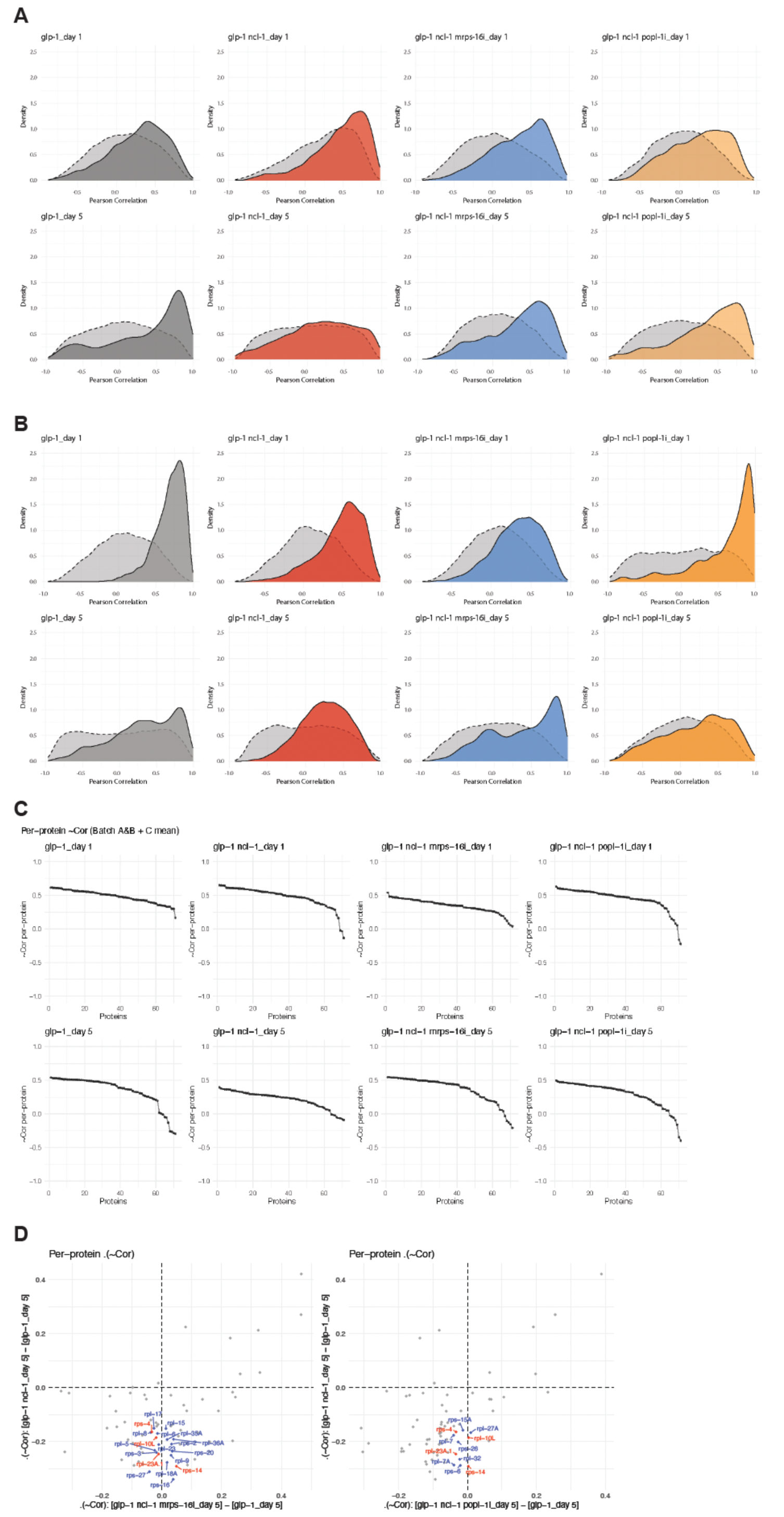
Loss of *ncl-1* leads to a loss of correlation between ribosomal proteins that can be restored by *mrps-16i* and *popl-1i*. B. Batch A distribution of pairwise protein-protein Pearson correlations among RPs at day 1 (top) and day 5 (bottom) in an experiment of single worm proteomics from *glp-1*, *glp-1 ncl-1*, *glp-1 ncl-1 mrps-16i*, and *glp-1 ncl-1 popl-1i* worm (∼15 worms per genotype). Dashed grey backgrounds indicate the background calculated with a set of proteins of matched abundance distribution to RPs. C. Batch B distribution of pairwise protein-protein Pearson correlations among RPs at day 1 (top) and day 5 (bottom) in a different experiment of single worm proteomics combined with *glp-1*, *glp-1 ncl-1*, *glp-1 ncl-1 mrps-16i*, and *glp-,1 ncl-1 popl-1i* worms (∼15 worms per condition per genotype). Dashed grey backgrounds indicate the background calculated with a set of proteins of matched abundance distribution to RPs. D. Per-protein average Pearson correlation from each RP to all other RPs at day 1 (top) and day 5 (bottom). Correlations were calculated from single worm proteomics data and combined across batches. Each point represents the mean correlation of a given ribosomal protein with all other RPs. E. Per-protein changes in average RP correlations in each genotype relative to *glp-1* at day 5. For each RP, the y-axis shows the change in mean Pearson correlation in *glp-1 ncl-1* relative to *glp-1* worms. The x-axis shows the changes in *glp-1 ncl-1 mrps-16i* vs *glp-1* (left panel) and *glp-1 ncl-1 popl-1i* vs *glp-1* (right panel). Blue dots indicate the RPs that show the strongest rescue towards glp-1 under the RNAi knockdowns, and red points indicate RPs commonly rescued by *mrps-16i* and *popl-1i*.

